# Aging and viral evolution impair immunity against dominant pan-coronavirus-reactive T cell epitope

**DOI:** 10.1101/2024.08.21.608923

**Authors:** Lucie Loyal, Karsten Jürchott, Ulf Reimer, Lil Meyer-Arndt, Larissa Henze, Norbert Mages, Jak Kostrzanowski, Bernhard Reus, Maike Mangold, Beate Kruse, Manuela Dingeldey, Birgit Sawitzki, Janine Michel, Marica Grossegesse, Karsten Schnatbaum, Holger Wenschuh, Andreas Nitsche, Nils Lachmann, Bernd Timmermann, Claudia Giesecke-Thiel, Julian Braun, Florian Kern, Andreas Thiel

**Affiliations:** Si-M / “Der Simulierte Mensch” a science framework of Technische Universität Berlin and Charité - Universitätsmedizin Berlin; Berlin, Germany; Berlin Institute of Health (BIH) at Charité – Universitätsmedizin Berlin, Immunomics - Regenerative Immunology and Aging; Berlin, Germany; JPT Peptide Technologies GmbH; Berlin, Germany; NeuroCure Clinical Research Center, Charité – Universitätsmedizin Berlin, corporate member of Freie Universität Berlin, Humboldt-Universität zu Berlin, and Berlin Institute of Health; Berlin, Germany; Department of Neurology with Experimental Neurology, Charité – Universitätsmedizin Berlin, corporate member of Freie Universität Berlin, Humboldt-Universität zu Berlin, and Berlin Institute of Health; Berlin, Germany; Max Planck Institute for Molecular Genetics; Berlin, Germany; Department of Informatics, School of Engineering and Informatics, University of Sussex; Brighton, UK; Department of Clinical and Experimental Medicine, Brighton and Sussex Medical School; Brighton, UK; Translational Immunology, Berlin Institute of Health (BIH); Berlin, Germany; Highly Pathogenic Viruses, Centre for Biological Threats and Special Pathogens, WHO Reference Laboratory for SARS-CoV-2 and WHO Collaborating Centre for Emerging Infections and Biological Threats, Robert Koch Institute; Berlin, Germany; Charité - Universitätsmedizin Berlin, Institute for Transfusion Medicine, Tissue Typing Laboratory; Berlin, Germany Berlin Institute of Health (BIH) at Charité – Universitätsmedizin Berlin, Immunomics - Regenerative Immunology and Aging; Berlin, Germany

## Abstract

Immune evasion by escape mutations subverts immunity against SARS-CoV-2. A role of pan-coronavirus immunity for more durable protection is being discussed but has remained understudied. We here investigated the effects of age, mutations, and homo-/heterologous vaccination regimens on the dominant pan-coronavirus-specific cellular and humoral epitope iCope after SARS-CoV-2 infection and vaccination in detail. In the older, quantitatively, and qualitatively reduced iCope-reactive CD4^+^ T cell responses with narrow TCR repertoires could not be enhanced by vaccination and were further compromised by emerging spike mutations. In contrast pan-coronavirus-reactive humoral immunity was affected only by mutations and not by age. Our results reveal a distinct deficiency of the dichotomous layer of pan-coronavirus immunity in the older, critical for long-term protection against SARS-CoV-2 variants.

**One-Sentence Summary:** Aging and viral evolution impair dominant pan-coronavirus immunity, a hallmark of efficient and broad immune competence against SARS-CoV-2

---

The severity of coronavirus disease 2019 (COVID-19) caused by severe acute respiratory syndrome coronavirus 2 (SARS-CoV-2) depends on a variety of factors including socioeconomic status, genetics, comorbidities, medication (immunosuppressants), pre-existing cross-reactive immunity and in particular on age (*1–5*). The rate of severe complications requiring hospitalization is low in the younger but increases beginning at the age of 30 resulting in a high fatality rate among the >80-year-olds (*6*). First generation vaccines provided effective protection to the Wuhan-WT SARS-CoV-2, however, the generally limited vaccine effectiveness in the older remains a concern (*7–9*). In addition, SARS-CoV-2 is undergoing rapid viral evolution generating variants of concern (VOC) characterized by increased transmissibility, altered disease severity, and evasion of neutralization by antibodies induced during previous vaccination and infection (*10–12*). Moreover, in general a rapid decline in mucosal antibodies providing direct protection at viral entry sites has been observed (*13*). These observations have dashed the hope that SARS-CoV-2 vaccines would be able to induce long-term, sterile humoral immunity. It is now rather emphasized that specific memory T cells provide a main layer for long-term protection, since they can react even against highly mutated variants due to their broad epitope coverage (*14–17*), including dominant pan-coronavirus epitopes (*18–20*). Such pan-coronavirus-reactive T cells lead to rapid cellular and humoral responses in the early phase of SARS-CoV-2 infection, which positively correlate with faster and better T cell and antibody responses after infection and vaccination (*4*, *5*, *8*, *18*, *21–24*). This observation links the presence of pre-existing pan-coronavirus-specific T cells directly to the success of early antibody induction. However, with age, spike-specific pan-coronavirus cellular immunity declines in older individuals (*8*, *18*, *25*, *26*). Whether this apparently critical component of cellular immunity can be restored in the older and how virus evolution affects cellular immunity directed at conserved spike protein regions is unknown. Importantly, cellular and humoral pan-coronavirus reactivity is dominated by an immunodominant coronavirus peptide (iCope) sequence located within the fusion peptide domain of the spike protein (amino acids 816-830) (*18*, *19*, *27*, *27–32*). We here comprehensively investigated the extent to which existing and potential future spike mutations impact iCope-specific cellular and humoral responses in unexposed individuals of different ages upon infection and vaccination.

## Results

### The iCope sequence is conserved in SARS-CoV-2 variants

First, we aimed to understand if and to what extent the SARS-CoV-2 spike S816-830 sequence SFIEDLLFNKVTLAD (iCope) is conserved. Therefore, we assessed the mutation rate of iCope in comparison to other sequences within spike. Within 10.8 mio globally reported spike sequence reads with unique mutations (GISAID, May 2022), the number of mutations per amino acid was comparatively low in iCope (Fig. 1A). Similarly, the summarized mutation rates in a scan of 15mers with 1 amino acid shifting revealed that iCope is a very inert sequence, especially compared to the highly mutated RBD sequence (aa 333-526) (Fig. 1B). In line with this, none of the existing variants of concern (VOC) from alpha to omicron including the omicron subvariants BA1-5 showed mutations within iCope. A comparison of iCope with the respective sequence of endemic coronaviruses (NL63, 229E, OC43, HKU1) demonstrated high conservation of the amino acids S816, E819, D820, L822, F823 and K825 (Fig. 1C). Only four out of 15 positions (N824, T827, L828, A829) displayed variation in the amino acid groups’ chemical property, while at positions F817, I818, L821 conservative substitutions retain the hydrophobic characteristics of the amino acids. Of 2.3 mio reads in the GISAD database by September 2021, 8 mutations affecting iCope with different incidences were reported (Fig. 1D). Consequently, to investigate potential alterations in T cell reactivity, we generated peptides for the identified mutations including the frequent hydrophobic-to-hydrophobic mutations: L822F, I818V, F817L and V826L. While the first mentioned three mutations are novel mutations, i.e. only occurring in SARS-CoV-2, the V826L mutation is shared by several HCoVs. The mutation T827A exchanges the polar T for the nonpolar A, whereas T827V, where T is exchanged for the equally hydrophobic V is also found in other HCoVs. D820N generates a novel mutation affecting an otherwise highly conserved amino acid. In order to also assess the effects of some drastic hypothetical changes in the amino acid sequence including changes to most conserved regions S816, E819, L821, F823, and K825 we additionally generated peptides with the following hypothetical mutations: S816D: polar to acidic, E819F: acidic to hydrophobic, L821K hydrophobic to basic, F823T: hydrophobic to polar and K825A: basic to nonpolar, N824L: polar to hydrophobic and V826D: hydrophobic to acidic (Fig. 1D).

**Fig 1:**
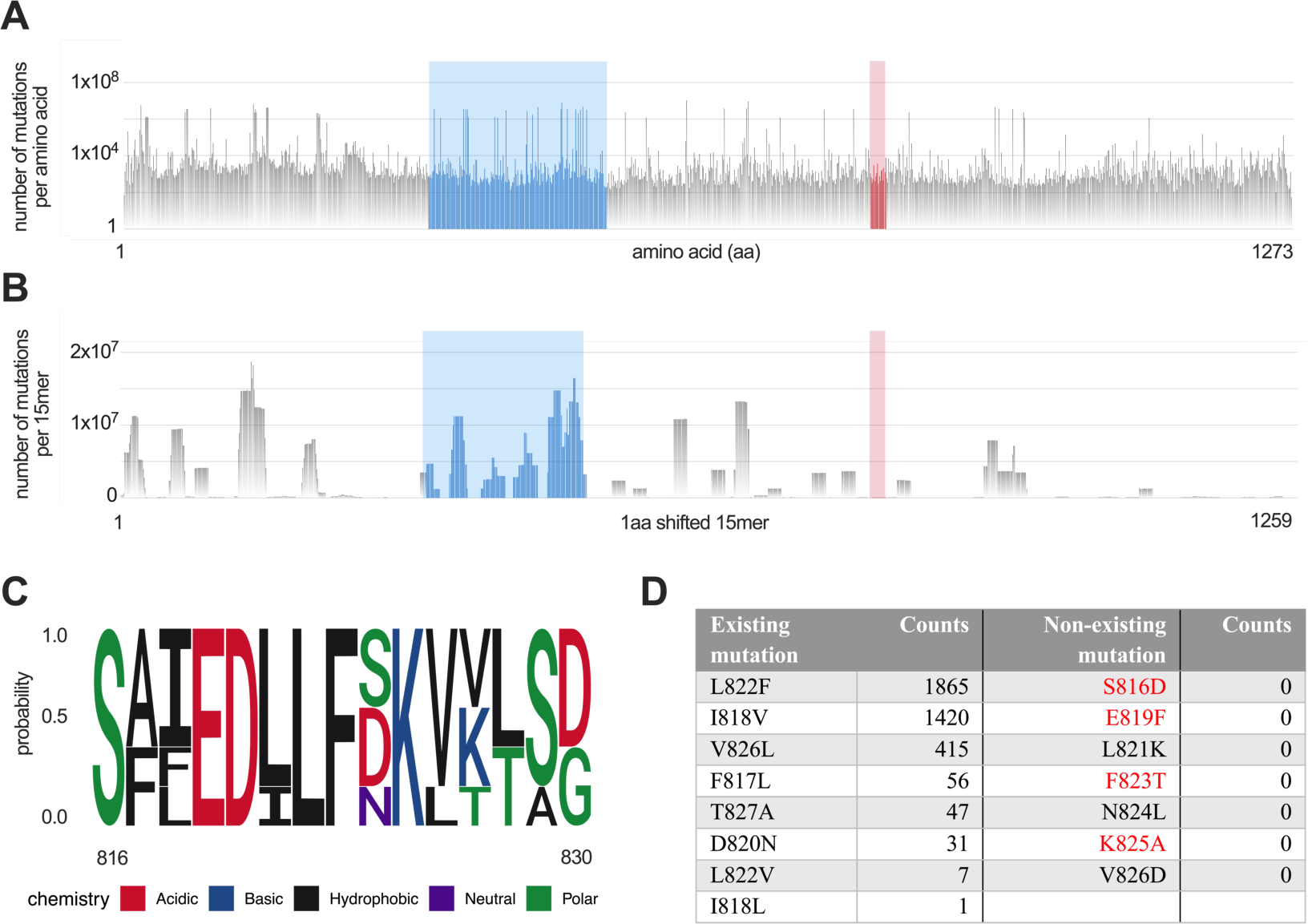
iCope mutation rate and sequence homology. **(A)** Mutations in 10.8 mio spike sequences per amino acid (GISAID, May 15^th^ 2022). Amino acids 816-830 (highlighted in red) compared to other amino acids within spike including RBD binding domain covering aa 333-526 (highlighted in blue). **(B)** Numbers of 1aa shifting 15mers carrying a mutation throughout spike (GISAID, May 15^th^ 2022). 15mers covering the iCope sequence are indicated in red. **(C)** Motif analysis of amino acid conservation of SARS-CoV-2 S816-830 in endemic coronaviruses (229E, NL63, OC43, HKU1). **(D)** Number of sequences with indicated mutations which were reported in GISAID by September 2021 and additionally generated non-existing mutations. Those replacing the most conserved amino acids are highlighted in red.

### The amino acids S819-826 are critical for iCope-specific T cell responses

We examined the effects of described and potential amino acid mutations on iCope-specific CD4^+^ T cell reactivity by comparing peptide stimulations of wildtype (WT) iCope to the different mutated iCope peptides as well as N-terminal (S-I, aa 1-643) and C-terminal (S-II, aa 633-1273) spike peptide pools. In 44% of young, unvaccinated and uninfected donors (unexposed, Table S1), we detected a strong response to stimulation with WT-iCope (Fig. 2A and B), and only a limited number of iCope mutations affected T cell responsiveness. Despite the high conservation of S816 across coronaviruses, the substitution of the polar S by the acidic D had no effect on T cell reactivity. While the V826D substitution impaired T cell activation, the conserved V826L mutation did not. The widely prevalent I818V mutation also had no effect on T cell reactivity. In contrast, the F817L mutation caused a decrease in 4 donors indicating a strong HLA-dependent effect. Strongest decrease of T cell reactivity was observed for mutations within the region S819- 826 (Fig. 2B and D) and thus, except for aa 824, the S819-826 region appears highly critical for effective T cell activation. Interestingly, recovery from COVID-19 does not appear to substantially increase the abundance of iCope-reactive CD4^+^ T cells (Fig. 2C), yet these cells have been shown to be engaged early during infection and vaccination with beneficial effects (*18*). In convalescent donors the effects of the different mutations follow the same pattern as in unexposed donors (Fig. 2D). Analysis of MHC class II /peptide binding prediction (IEDBD.org) in combination with iCope-specific T cell responses comprising all 96 donors of Table S1 indicated that the mutations S816D, F817L, I818L, I818V, N824L, V826L, T827A did not significantly affect peptide binding and T cell responsiveness (Fig. 2E and Fig. S1). While L821K, F823T and V826D were identified in silico as poor MHC binders (depicted in light grey, Fig. 2E) also the mutations E819F, D820N, L822F, L822V and K825A impair the anti-iCope response (depicted in dark grey, Fig. 2E). Altogether, this identifies the S819-826 region as critical for iCope-specific T cell responses.

**Fig 2:**
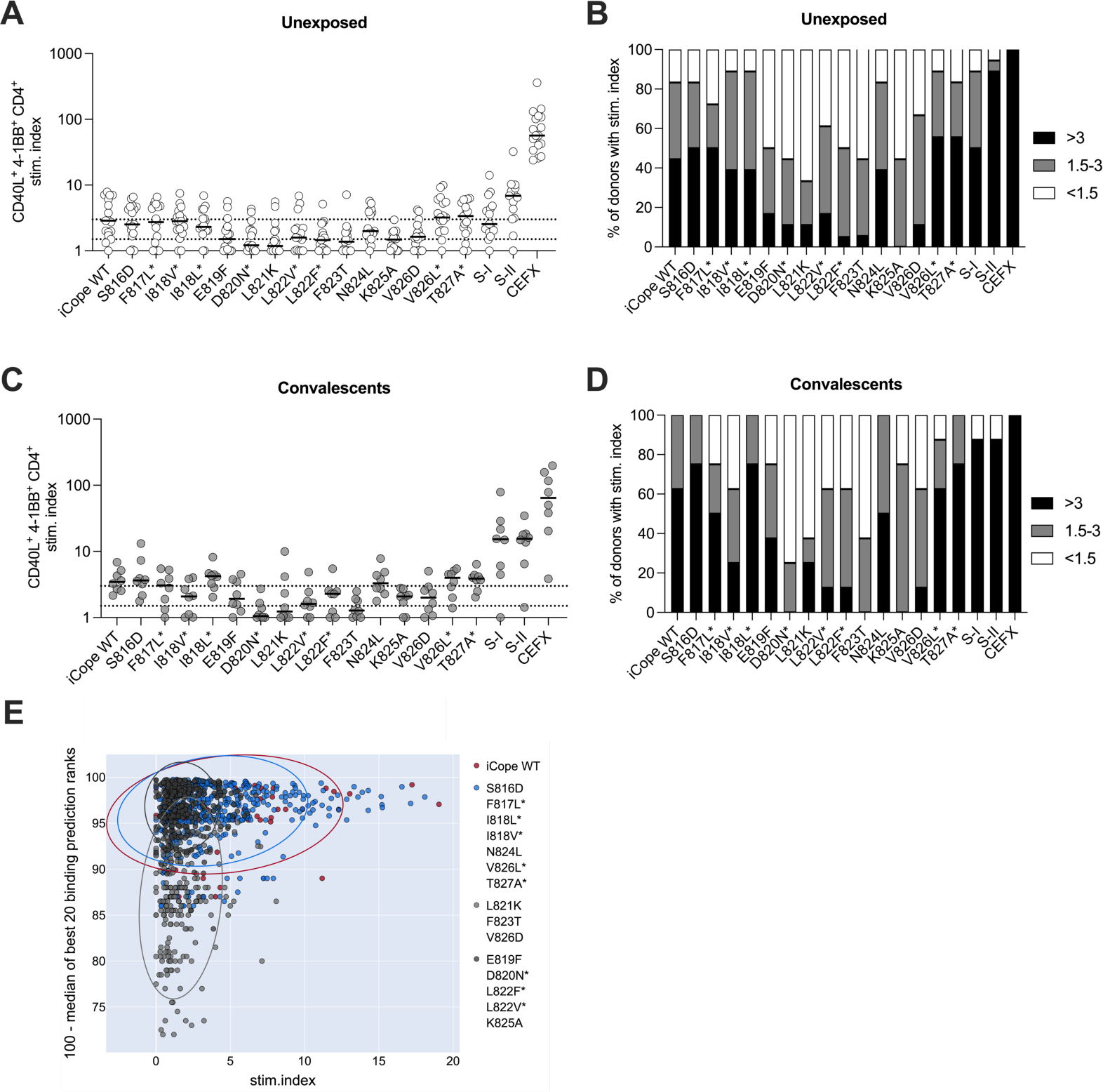
Mutations in S819-826 impair iCope specific T cell responsiveness in unexposed and convalescents. Ex vivo stimulation of PBMCs from unexposed individuals (*n*=17) and COVID-19 convalescents (*n*=8) with iCope WT or different mutated iCope peptides and control pools S-I and S-II, and CEFX as positive control. **(A, C)** The percentage of CD40L^+^4-1BB^+^ CD4^+^ T cells among stimulated PBMC was divided by the percentage of these cells among unstimulated PBMC to determine the stimulation index (stim. index) shown on the y-axis. Dotted lines indicate a stim. index of 1.5 and 3. All values below 0.1 were set at 0.1 for display. (**B, D)** Bar plots show the proportions of individuals with the indicated peptide or peptide pool stimulations with a stim. Index >3 (black), 1.5-3 (grey) or <1.5 (white). **(E)** MHCII peptide binding was predicted for all combinations of the iCope/mutations peptides for all 96 donors from Table S1 using IEDB. The median of the best 20 prediction ranks was calculated for each allele and plotted against the stimulation index from the stimulations with iCope or its mutant peptides. Color codes indicate iCope wildtype peptide (red), mutated iCope peptides with no to weak negative effect on T cell activation (blue) and mutated peptides with strong negative effect on T cell activation (grey). Those with IEDB based, predicted negative effect are separated from the others by light grey dots. * indicates documented iCope mutations.

### Vaccination boosts iCope immunity in the younger but not in the older

We have previously shown that vaccination with the BNT162b2 (BNT, BioNTech/Pfizer) mRNA COVID-19 vaccine leads to activation of iCope-specific T cells with beneficial effects. However, the level of pre-existing S-II spike peptide pool-specific cross-reactive T cells, dominated by iCope-specific cells, decreased with age (*18*). To address the consequences of this on vaccine immunogenicity in the older, we compared the CD4^+^ T cell responses to iCope upon homologous BNT162b2 vaccination in young (age <40, mean 30.8 years) and older individuals (age >60, mean 76.5 years, Table S1). While the impact of mutations remained comparable in both groups, the total amount of responsive T cells in the older, triple vaccinated donors decreased to a level similar to that of young, unexposed donors (Fig. 3A, B, cf. Fig. 2A). When we compared the effects of the different vaccination regimens, we found that T cell responsiveness was not affected by vaccine combination, but again by age (Fig. 3C-E). While vaccination enhanced iCope-specific cellular responses in the younger and also to potential mutants (cf. Fig. 2A and 3A), neither homologous nor heterologous vaccination could rescue the low iCope-specific CD4^+^ T cell responses in older individuals (Fig. 3E).

**Fig 3:**
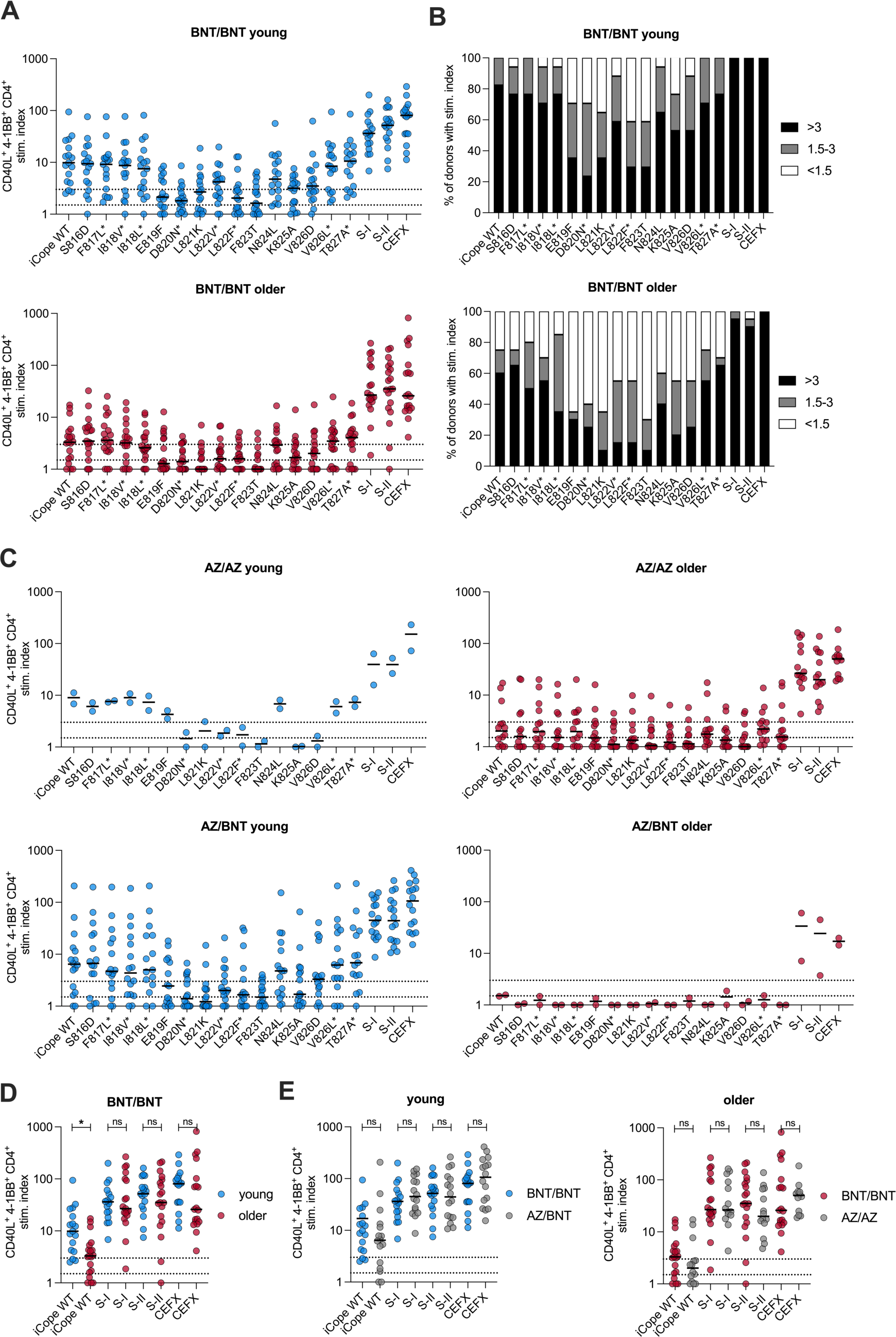
iCope responsiveness in vaccinated donors is age dependent. **(A-E)** Ex vivo stimulation of PBMCs from young or older BNT/BNT (*n*=18/19), AZ/AZ (*n*=2/14), and AZ/BNT (*n*=16/2) vaccinated individuals with iCope WT or different mutated iCope peptides and control pools S-I, S-II and CEFX. * indicates documented iCope mutations. **(A, C, D, E)** The percentage of CD40L^+^4-1BB^+^ CD4^+^ T cells among stimulated PBMC was divided by the percentage of these cells among unstimulated PBMC to determine the stimulation index (stim. index) shown on the y-axis. Dotted lines indicate a stim. index of 1.5 and 3. All values below 0.1 were set at 0.1 for display. **(B)** Bar plots show the proportions of individuals with the indicated peptide or peptide pool stimulations with a stim. index >3, 1.5-3 or <1.5. **(D)** Comparison of young and older BNT/BNT vaccinated donors under the indicated stimulation conditions. \**P*<0.05, \*\**P*<0.01, \*\*\**P*<0.001 and ns for *P*>0.05 (Student’s *t* test). **(E)** Comparison of young BNT/BNT with young AZ/BNT vaccinated donors and older BNT/BNT with older AZ/AZ vaccinated donors under the indicated stimulation conditions. \**P*<0.05, \*\**P*<0.01, \*\*\**P*<0.001 and ns for *P*>0.05 (Student’s *t* test). BNT: BNT162b2 mRNA COVID-19 vaccine (BioNTech); AZ: ChAdOx1 COVID-19 vaccine (AstraZeneca).

### Age and mutations affect the quality of the iCope specific T cell but not B cell response after infection and vaccination

Next, we assessed whether mutations in the iCope sequence and/or aging affect the quality of the iCope-specific T cell responses, i.e. TCR avidity and effector functions, upon infection or vaccination. We used the CD3 surface expression level among antigen-reactive CD4^+^ T cells as readout as the degree of CD3 downregulation (CD3^lo^) on *in vitro* activated T cells can serve as a surrogate for TCR avidity (*16*). The TCR avidity of the iCope-specific T cells was significantly reduced by mutations within the aa 819-823 region of spike as well as in position aa 825-826 in all examined cohorts (Fig. 4A and Fig. S2A). In general, peptides with mutations leading to reduced MHC-binding also induced lesser CD3 downregulation, however there were some exceptions such as F817V and I818L with a negative impact on the TCR avidity but not overall T cell responsiveness (Fig. 4A and Fig. S2A). The L822V replacement reduced the amount of reactive cells but the few responding cells showed high avidity for the target. In the course of SARS-CoV-2 infection high avidity T cells targeting S-I and S-II peptides were generated but no substantial changes in the avidity of iCope-specific T cells was observed after clearance of the infection. In contrast to this, the homologous BNT prime-boost (BNT/BNT) resulted in the expansion of high avidity iCope-specific T cell clones in the young, but again not in the older individuals (Fig. 4A and B). Moreover, remaining iCope-specific T cells in vaccinated older people displayed a reduced capacity for TNF-α and IFN-γ secretion (Fig. 4C), two cytokines fundamental in the hosts combat to SARS-CoV-2 infection. Mutations, too, led to an impaired polyfunctional response in the remaining iCope-specific T cells (Fig. S2B). In contrast, anti-iCope antibodies were increased upon infection and vaccination along with the SARS-CoV-2 S1- and S2-specific humoral immunity with no detectable difference between the young and the older vaccinees (Fig. 4D and E). When assessing the impact of different mutations to antibody binding, F823T together with S816D, I818V, and E819F displayed the strongest negative effect on antibody binding, whereas for T cells any tested mutation affecting the region between E819F-V826D had a strong negative effect (Fig. S3A). In summary, aging drastically affects the composition and functional capacity of iCope-specific T cells but not humoral responses. Mutations however seem to affect both cellular and humoral immunity albeit to different extents.

**Fig 4:**
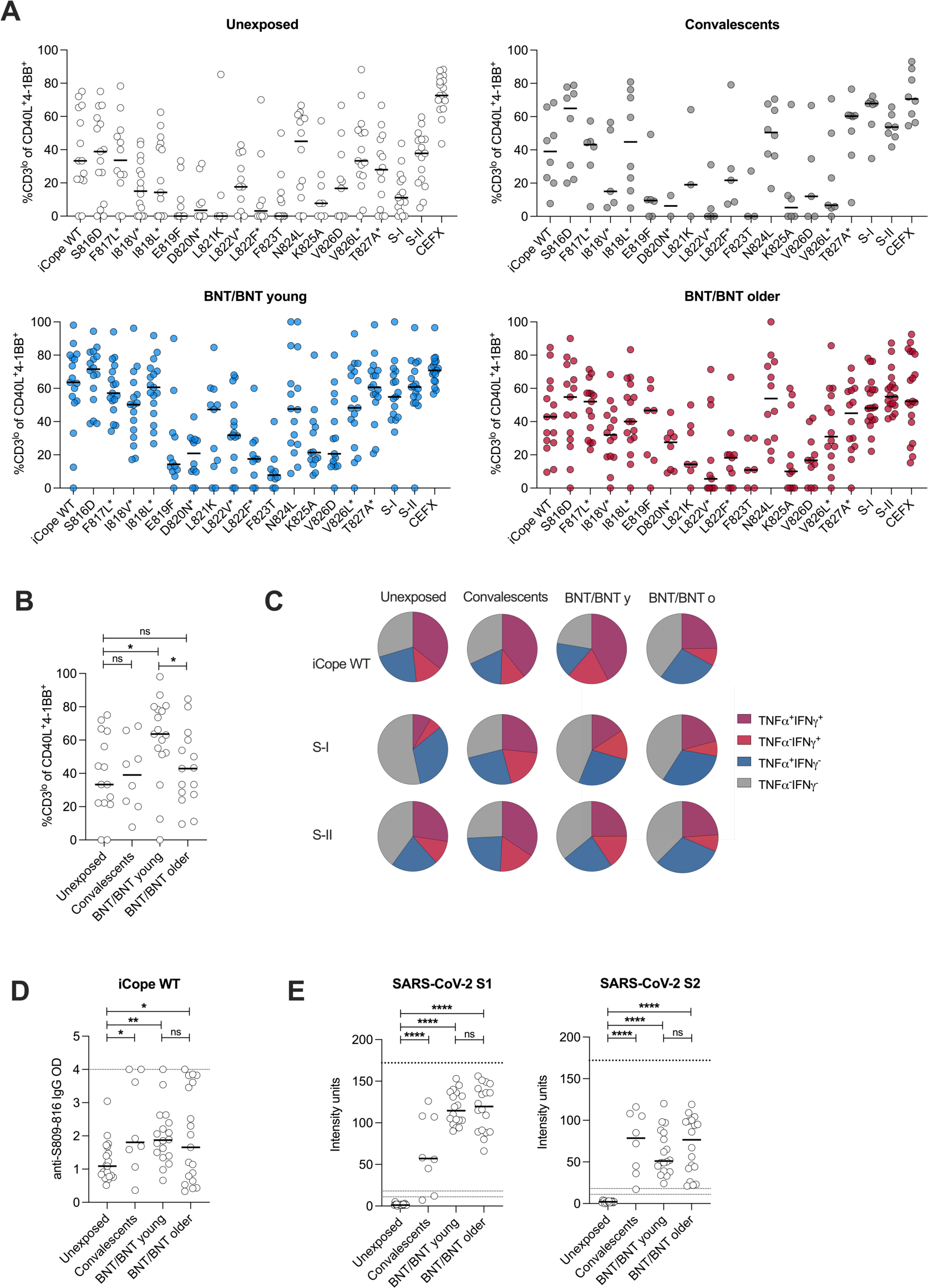
The quality of iCope responsiveness is impaired in older individuals. Ex vivo stimulation of PBMCs from unexposed (*n*=17), convalescent (*n*=8), young BNT/BNT vaccinated (*n*=18) and older BNT/BNT vaccinated (*n*=19) individuals with iCope WT or different mutated iCope peptides and the control pools S-I, S-II and CEFX. * indicates documented iCope mutations. **(A, B)** Frequencies of CD3^lo^ cells in CD40L^+^4-1BB^+^ CD4^+^ T cells. CD3^lo^ frequencies are shown for T cell responses with a stim. index ≥1.5. **(C)** Proportion of IFN-γ and/or TNF-α producing T cells among CD40L^+^4-1BB^+^ CD4^+^ T cells in the indicated cohorts after stimulation with iCope WT, S-I or S-II. **(D)** Optical density (OD) of anti-S809-826 wildtype (WT) peptide IgG (ELISA) in unexposed (*n*=17), convalescents (*n*=8), BNT/BNT young (*n*=18) and older BNT/BNT (*n*=19). **(E)** Levels of anti-S1- or anti-S2-IgG binding antibody intensity units in indicated cohorts. Dotted lines indicate lower cut-off (set at 11) for definitive negative values and a range from 11-18 for values classified as positive with uncertainty. \**P*<0.05, \*\**P*<0.01, \*\*\**P*<0.001, \*\*\*\**P*<0.0001 and ns for *P*>0.05 (Student’s *t* test).

### Reduced iCope T cell responsiveness in the older is associated by a thinned-out TCR repertoire

To delineate the age-associated quantitative and qualitative differences of iCope-specific T cells in more detail and determine whether additional vaccinations could rescue the observed scrambled responses in the older, we assessed iCope and S-II responses in individuals aged <40 or >70 who had three vaccinations but had no indicators of infection (anti-nucleocapsid IgG titer and/or anti-nucleocapsid T cell response) (Fig. 5A and B; Fig. S4A and B). We observed robust (>200 days) T cell frequencies against S-II in the older (Fig. 5A and Fig. S4A) however, the number of donors that were weak- to non-responsive against the pan-coronavirus-specific iCope remained high even after the third vaccine dose (Fig. 4B, S4B). To deconvolve the reason for the altered iCope-specific T cell responses in the older we conducted single-cell RNA-sequencing of FACS-purified iCope- reactive CD4^+^ T cells (CD40L^+^4-1BB^+^) and as a control iCope-non-reactive (CD40L^-^4-1BB^-^) CD4^+^ T cells from 3 young (<30 years) and 3 older (>80 years) donors 3 months after the third dose of BNT162b2 vaccine and 6 young and 5 older equally vaccinated donors with SARS-CoV- 2 infection in the last 3 months prior sampling (Table S2). iCope-non-reactive CD4^+^ T cells showed an age-related shift on molecular level whereas iCope-specific T cells showed no age-related cell-intrinsic gene expression differences except of a distinct population of cytotoxic CD4^+^ T cells the red cluster in the upper left (Fig. 5C and D). However, the overall and iCope-reactive TCR repertoire diversity was reduced in the older, characterized by an enrichment of just few clones (Fig. 5E, F). Assessing the proportion of iCope-specific TCR CDR3 sequences of CD40L^+^4-1BB^+^ within CD40L^-^4-1BB^-^ T cells we observed no differences between young and older donors arguing against an age dependent activation deficiency (Fig. 5G). Non-activated iCope-specific clones among CD40L^-^4-1BB^-^ T cells in older also did not display any sign of exhaustion, anergy or senescence but a prominent cytotoxic CD4^+^ T cell signature (Fig. 5H). Our data suggests that the reduction in quantity and quality of iCope-specific T cells in the older is caused by a narrowing of the TCR repertoire accompanied by a loss of clones with high TCR avidity and effector functions and differentiation into cytotoxic CD4^+^ T cells.

**Fig 5:**
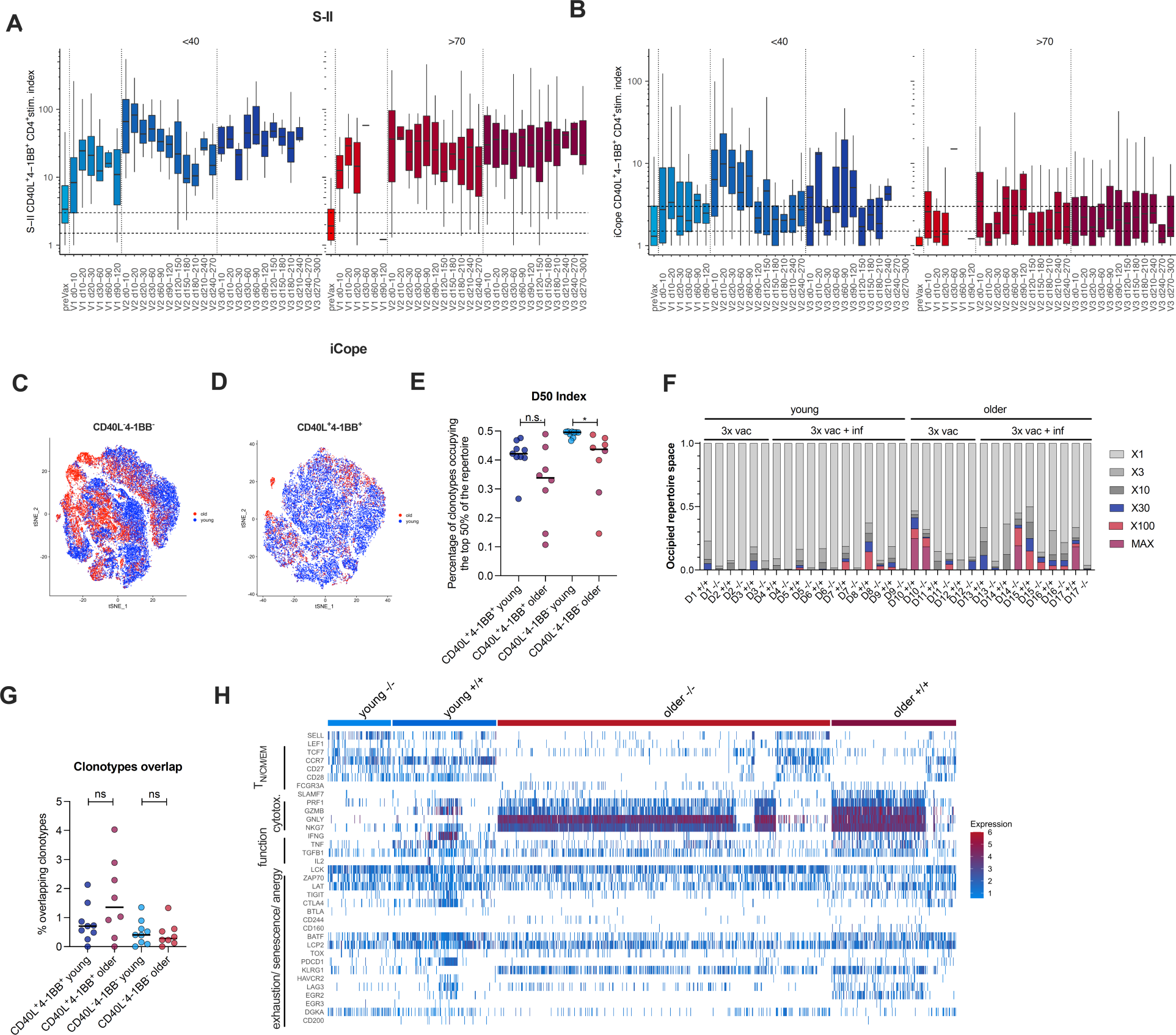
Older people possess functional iCope-reactive T cells but with limited clonal breadth. **(A, B)** Ex vivo stimulation of PBMCs from young (<40 years, blue, *n*=507) or older (>70 years, red, *n*=1267) individuals with S-II **(A)** or iCope **(B)** prior to first vaccination (preVax) or at indicated times after the first, second or third dose of vaccination (V1-3). The percentage of CD40L^+^4-1BB^+^ CD4^+^ T cells among stimulated PBMC was divided by the percentage of these cells among unstimulated PBMC to determine the stimulation index (stim. index) shown on the y-axis. **(C, D)** Single-cell gene expression of FACS-purified iCope reactive CD40L^+^4-1BB^+^ and CD40L^-^4-1BB^-^ CD4^+^ T cells of *n*=9 young (<43 years) and *n*=8 older (>76 years) donors. tSNE visualization of CD40L^+^4-1BB^+^ **(C)** and CD40L^-^4-1BB^-^ **(D)** CD4^+^ T cell clustering of young (blue dots) and older (red dots) donors. **(E)** D50 diversity index indicating the number of clonotypes occupying the top 50% of the TCR repertoire. **(F)** Proportion of clonotypes with specific counts in indicated samples of the different donors (D1-6). D1-3 are the triple vaccinate young donors without, D4-9 with recent infection, D10-12 triple vaccinated older donors without, 13-17 with recent infection. **(G)** Frequencies of overlapping clonotypes between activated CD40L^+^4-1BB^+^ and inactivated CD40L^-^4-1BB^-^ CD4^+^ T cells in young and older donors. **(H)** Heatmap of gene expression signatures of curated gene sets (as given in the Material and Methods section) for naïve and effector T cell characteristics as well as cytotoxicity, anergy, exhaustion and senescence in overlapping clonotypes between activated CD40L^+^4-1BB^+^ (+/+) and inactivated CD40L^-^4-1BB^-^ (-/-) CD4^+^ T cells.

## Discussion

The future dynamics and socioeconomic impact of SARS-CoV-2 and potential future coronavirus spillovers strongly relies on several factors including the extent and effect of existing immunity. This immunity varies due to pre-existing reactivities, vaccinations and encountered infections, and individual susceptibility to severe disease, which is influenced by age, comorbidities, medications (e.g. immunosuppression), and genetics. Newly arising variants of concern can differ drastically with respect to their immunogenicity and hence the immune system’s ability to recognize and neutralize them at sites of viral entry. The so far reported mutations in the main antigenic determinant of SARS-CoV-2, the spike glycoprotein, predominantly occur in the N-terminal and receptor-binding domains (NTD and RBD) and overcome sterile, antibody mediated immunity at the mucosal surface (*11*). The currently latest VOC, Omicron, has accumulated sufficient numbers of critical mutations to eliminate the effect of most of the neutralizing antibodies in vaccinated but also convalescent individuals (*12*).

Recurrent booster vaccinations appear necessary to maintain effective albeit low levels of neutralizing antibodies providing protection (*33*, *34*). In contrast, both, infection and vaccination, induce broad specific T cell repertoires largely retaining intact Omicron-specific T cell responses in both convalescents and vaccinated individuals (*14*, *35*). Due to the MHC polymorphism mutations may not only result in the loss of selected individual T cell epitopes but also generate new ones (*36*). Interestingly, however, not all combinations of vaccinations and infections with SARS-CoV-2 variants boost T cell responses. Hybrid immunity resulting from infection with Wuhan-WT SARS-CoV-2 followed by an Omicron infection results in "hybrid immunity dampening" with reduced T cell responses against the S1-subunit (*36*). Importantly, the effect is not observable assessing S2-subunit-specific T cell responses, suggesting the largely unmutated S2 epitopes of spike can compensate for this effect.

The second most fundamental challenge for efficient long-lasting immunity against SARS-CoV-2 and its emerging variants is the aging of the human immune system that is associated with a loss of a broad polyclonal repertoire of naïve T cells caused by thymic involution beginning with puberty (*37–40*). Reduced thymic output has been demonstrated to affect primary vaccination in the older even with live vaccines such as yellow fewer vaccine (*41*, *42*). Low numbers of naïve T cells and lower TCR diversity against SARS-CoV-2 epitopes were also associated with severe COVID-19 (*43*, *44*) and lower frequencies of IFN-γ- and IL-2-producing T cells in SARS-CoV-2 have been reported in vaccinated older (*45*). Accordingly, pre-existing pan-coronavirus-specific memory T cells may be a critical source of cellular immunity to enable timely reactivation of pre-existing memory B cells but also for the fast recruitment of new naïve B cell specificities (*46*, *47*). Pre-existing cross-reactive S2-specific antibodies can prevent infection by blocking the fusion machinery of SARS-CoV-2 and IgA producing B cells have been demonstrated already after the first dose of vaccination (*27*, *48–55*). Given the possible importance of neutralizing S2-specific antibodies for controlling SARS-CoV-2 variants characterized by altered RBD regions, the evolution of these responses should be analyzed in more detail. Additionally, the mechanisms how these responses are maintained also in the older, should be explored. It is of interest in this context that in influenza infection rapid cellular responses mediated by pre-existing cross-reactive T cells are associated with protection from symptomatic infection and severe disease even in seronegative individuals (*56–58*). In line with this, pre-existing cross-reactive T cells are able to provide T cell help in the early phase of infection or after vaccination and protect against SARS-CoV-2 infection (*4*, *18*, *21*). Additionally, pre-existing cross-reactive T cells targeting the SARS-CoV-2 polymerase can abort infection and so contribute to the containment of virus spreading (*5*).

The pan-coronavirus-specific fusion peptide region in spike contains antigenic determinants recognized by B cells and T cells that so far have remained relatively unaffected by novel mutations of SARS-CoV-2. This region may, therefore, be considered as natural target for the development of pan-coronavirus specific vaccines. However, pre-existing pan-coronavirus-specific cellular immunity declines with age and we reveal here that the resulting deficiencies cannot be restored by homologous or heterologous vaccination in older individuals. While in young individuals a broad and diversified iCope-specific clonotype repertoire is induced by vaccinations or in combination with infections older people are not capable of such repertoire broadening. Our results suggest that the impaired pan-coronavirus-specific T cell responsiveness to SARS-CoV-2 vaccination is caused by gradual loss of "best-fit" clones, an effect also known from influenza (*59*, *60*). This is also in line with the recent observation that cross-reactive responses are mainly established in early childhood and decline afterwards (*26*). Additionally, remaining responses in the older appear to be characterized by a low TCR avidity towards iCope, which can result in the observed reduced polyfunctionality. Besides, we did not observe signatures of senescence, exhaustion or anergy in iCope-specific clones (*61*). Even after a third vaccination the lack of reactive clones in the repertoire rather than unresponsiveness of existing clones (e.g. due to anergy or exhaustion) is the reason for impaired T cell responsiveness in the older. It has been emphasized that in many older individuals a third dose of vaccine is necessary to boost an incomplete response against the complete spike protein achieved with two vaccinations (*7*). However, we reveal here that even a booster vaccination may result in, albeit quantitatively equal, qualitatively still impaired responses.

Single-cell RNA sequencing revealed expanded iCope specific cytotoxic CD4^+^ T cell population in the older donors, which were partly not identified as antigen-specific due to the absent CD40L expression in terminally differentiated cytotoxic CD4^+^ T cells (*62*). Accumulation of cytotoxic CD4^+^ T cells is a general feature of the very old but their enrich has been observed also in adults compared to children and hospitalized patients compared to non-hospitalized individuals (*63*, *64*). Strong cytotoxic CD4^+^ T cell responses early in the infection were linked to defects in B cell responses implying iCope- and therefore pan-coronavirus specific T cells with cytotoxic properties might be a risk factor for severe disease (*65*).

The development of next generation pan-coronavirus-targeting vaccines focusing on intranasal administration could compensate for the observed alterations in the older, if sufficient reservoirs of tissue resident T cells are still present that can be re-activated by intranasal challenge. It has been already demonstrated that high titers of neutralizing antibodies can be induced in nasal administrated ChAdOx1 in hamsters (*66*) and in phase I clinical trials with inactivated NDV-HXP-S vaccines (*67*). Other first results are discouraging, revealing rather low induction of specific immunity (*68*). Utilizing dominant anti-SARS-CoV-2 epitopes derived from different SARS-CoV- 2 proteins a strong T cell induction was observed, however the stability of these responses remain to be assessed (*69*). Previous and recurrent encounters with endemic coronaviruses and more recently with different SARS-CoV-2 variants results in high levels of pan-coronavirus reactive T- and B-cells in the respiratory tract as shown recently for bronchoalveolar lavage (BAL) and oropharyngeal lymphoid tissue (*70*, *71*). We propose that only with the efficient local (re-)activation of these pan-coronavirus-reactive immune cells long-term immunity against SARS-CoV-2 and its emerging variants is achievable. In particular in the elderly this reservoir might be the only source of sufficiently broadly targeting immune cells against SARS-CoV-2, since the availability of broadly specific naïve T cells in the older is irrevocably decreased.

## Materials and Methods

### Study participants

This study was approved by the Institutional Review board of the Charité (EA/152/20). Written informed consent was obtained from all included participants and the study was conducted in agreement with the declaration of Helsinki. Age and gender of all donors was recorded (Table S1). Participants who had tested positive for SARS-CoV-2 RNA (RT-qPCR from nasopharyngeal swabs) and displayed SARS-CoV-2 related symptoms were classified as convalescent donors. All convalescent donors had mild symptoms (WHO severity grade < 3) and no hospitalization was required. The day of infection was set as day –3 prior to reported symptom onset. The vaccinated cohorts had following intervals between their first and second dose of vaccine: homologous BNT162b2 vaccination: 3 weeks, homologous ChAdOx1 vaccination: 12 weeks and heterologous ChAdOx1/BNT162b2 vaccination: 12 weeks. All enrolled participants were screened for SARS-CoV-2 subunit 1 (S1) and nucleocapsid (N) antibody titers. Two unexposed, two young BNT/BNT vaccinated and one older BNT/BNT vaccinated individual were excluded from the analysis due to detectable S1 and/or N (unexposed) or N (vaccinated) IgG titers.

### Coronavirus RT-qPCR

RNA was extracted from 140 μl of wet nasopharyngeal swabs (Copan mini UTM) using the QIAamp Viral RNA Mini Kit and QIAcube Connect with the manual lysis protocol. SARS-CoV- 2 RNA detection was performed using a simultaneous two duplex one-step real-time RT-PCR assay with custom primers and probes (Metabion and Thermo Fisher Scientific) for SARS-CoV-2 E Gene and SARS-CoV-2 ORF1ab according to the RKI/ZBS1 SARS-CoV-2 protocol as described before (*72*). Each one is duplexed with a control that either indicates potential PCR inhibition or proves the successful extraction of nucleic acid from the clinical specimen. As positive controls genomic SARS-CoV-2 RNA and genomic SARS-CoV RNA were used for the ORF1ab and the E-Gene assay, respectively, adjusted to the Ct values 28 and 32. PCR was conducted with the AgPath-ID™ One-Step RT-PCR Reagents kit (Applied Biosystems) using a Bio-Rad CFX96 or Bio-Rad Opus real-time PCR cycler.

### Blood and serum sampling and PBMC isolation

Whole blood was collected in lithium heparin tubes for peripheral blood mononuclear cells (PBMC) isolation and SST™II advance (all Vacutainer^®^, BD) tubes for serology. SST™II advance tubes were centrifuged for 10min at 1000g prior to removing serum. Serum aliquots were frozen at –20°C until further use. PBMCs were isolated by gradient density centrifugation according to the manufacturer’s instructions (Leucosep tubes, Greiner; Biocoll, Bio&SELL).

### SARS-CoV-2 IgG S1 and N ELISA

Anti-SARS-CoV-2 IgG ELISA specific for spike subunit 1 (S1) and nucleocapsid (N) were performed with a 1:100 serum dilution using the commercial kits (EUROIMMUN Medizinische Labordiagnostika AG) according to the manufacturer’s instructions and measured at a Tecan infinite M plex reader with Magellan Pro V7.4 software. The test results were considered positive with uncertainty within an OD ratio (defined as absorbance difference between control and study sample) of 0.8-1.1 and positive above >1.1

### Anti-IgG Immunoblot

Multiplex anti-IgG Immunoblot specific for the spike subunit 1 (S1), spike subunit 2 (S2), was performed using the EUROLINE Anti-SARS-CoV-2-Profil (IgG) kit (EUROIMMUN Medizinische Labordiagnostika AG) according to the manufacturer’s instructions. The data was analyzed using the EUROLine Scan Software. Lower cut-off was set at 11 for certainly negative values. The range from 11-18 indicates the values classified as positive with uncertainty.

### Epitope-specific antibody ELISA

400 nM of biotinylated peptide S809-826 (Biotin-Ttds-PSKPSKR*SFIEDLLFNKV*-OH (Ttds linker=N-(3-{2-[2-(3-Amino-propoxy)-ethoxy]-ethoxy}-propyl)-succinamic acid, JPT Peptide Technologies) or mutated peptides as indicated in the figure legends were immobilized on a 96 well Streptavidin plate (Steffens Biotechnische Analysen GmbH) for 1 hour at RT. After blocking (1 hour, 30°C) serum samples were diluted 1:100 and incubated for 1 hour at 30°C. HRP-coupled, anti-human-IgG secondary antibody (Jackson Immunoresearch) was diluted 1:5000 (Jackson Immunoresearch) and added to the serum samples for 1 hour at 30°C, then HRP substrate was added (TMB, Kem-En-Tec). The reaction was stopped by adding sulfuric acid and absorption was measured at 450 nm using a FlexStation 3.

### Ex vivo T cell stimulation

Freshly isolated PBMC were cultivated at a concentration of 5*10^6^ PBMC/ml in AB-medium containing RPMI 1640 medium (Gibco) supplemented with 10% heat inactivated AB serum (Pan Biotech), 100 U/ml of penicillin (Biochrom), and 0.1 mg/ml of streptomycin (Biochrom). Stimulations were conducted with PepMix^TM^ overlapping peptide pools (15 aa length with 11 aa overlaps, JPT Peptide Technologies) covering the N-terminal (S-I) and C-terminal (S-II) part of spike glycoprotein. Single peptide stimulations were conducted with the following peptides: iCope WT (Ń-SFIEDLLFNKVTLAD-Ć) or the mutated iCope peptides with following aa substitutions: S816D, F817L, I181V, I818L, E819F, D820N, L821K, L822V, L822F, F823T, N824L, K825A, V826D, V826L, T827A (all JPT Peptide Technologies). All stimulations (peptide pools and single peptides) were performed at final concentrations of 1 µg/ml per peptide. For negative control the stimulation peptide solvent DMSO diluted 1:1 in PBS was used at the same concentration as in peptide-stimulated tubes. CEFX Ultra SuperStim pool (1 µg/ml per peptide) (JPT Peptide Technologies) were used as positive stimulation controls. For optimized costimulation, purified anti-CD28 (clone CD28.2, BD Biosciences) was added to each stimulation at a final concentration of 1 µg/ml. Incubation was performed at 37°C, 5% CO_2_ for 16 hours in the presence of 10 µg/ml brefeldin A (Sigma-Aldrich) during the last 14 hours. CD4^+^ T cell activation was calculated as a stimulation index (stim. index) = % of CD40L^+^4-1BB^+^ CD4^+^ T cells in the stimulation / % of CD40L^+^4-1BB^+^ CD4^+^ T cells in the unstimulated control. Dotted lines indicate a stim.index of 1.5 (positive with uncertainty) and 3 (definite positive). All values below 0.1 were set as 0.1 for display.

### Flow Cytometry

Stimulations were stopped by incubation in 2 mM EDTA for 5 min. Surface staining was performed for 15 min in the presence of 1 mg/ml of Beriglobin (CSL Behring) with the following fluorochrome-conjugated antibodies titrated to their optimal concentrations: anti-CD3-FITC (Miltenyi Biotec), anti-CD4-VioGreen (Miltenyi Biotec), anti-CD8-VioBlue (Miltenyi Biotec), anti-CD38-APC (Miltenyi Biotec), and anti-HLA-DR-PerCpVio700 (Miltenyi Biotec). During the last 10 min of incubation, Zombie Yellow fixable viability staining (Biolegend) was added. Fixation and permeabilization were performed with eBioscience^TM^ FoxP3 fixation and PermBuffer (Invitrogen) according to the manufacturer’s protocol. Intracellular staining was carried out for 30 min in the dark at room temperature with anti-4-1BB-PE (Miltenyi Biotec), anti-CD40L- PEVio770 (Miltenyi Biotec), anti-IFN-γ (Biolegend) and anti-TNF-α V605 (Biolegend). All samples were measured on a MACSQuant^®^ Analyzer 16 (Miltenyi Biotec). Instrument performance was monitored prior to every measurement with Rainbow Calibration Particles (BD Biosciences).

### HLA typing and analysis

HLA typing was performed by LABType CWD assays (One Lambda, West Hills, CA, USA) based on reverse sequence-specific oligonucleotides (rSSO) according to the manufacturer’s instructions. Briefly, the HLA genomic region was amplified individually using locus-specific biotinylated primers for HLA-DRB1, -DQA1, -DQB1, -DPA1, and –DPB1. Amplicons were hybridized to HLA allele- and allele-group-specific probes attached to Luminex® beads. Complementary binding was detected by addition of R-phycoerythrin-conjugated streptavidin and acquired using a FLEXMAP 3D flow analyzer (Luminex, Austin, TX, USA). HLA alleles were derived at two-field code resolution (highest probability) as referenced in the catalogue of common and well-documented HLA alleles version 2.0.0 33. 12 individuals with a mean stimulation index across iCope/mutations outside 1.5 times the interquartile range from the upper or lower quartile were removed as outliers. MHCII peptide binding predictions were then generated for all combinations of the iCope/mutations peptides and alleles using the IEDB tools api (v2.26). All lengths from 11 to 15 were selected and the percentile rank was taken as the binding rank. Results where the allele was only typed to the first field (e.g., DPA1*02) and alleles with unavailable prediction methods were discarded. Each individual’s binding prediction results were collated to a list of all the results for their 2 DRB1 alleles, their 4 combinations of DQ, and their 4 combinations of DP. Each donor would then have 16 sets of collated binding results, one for each peptide. The median of the best 20 prediction ranks was calculated for each allele/peptide combination. Where 20 predictions were not available, all available predictions were used. For the MCHII- medianbest20ranks/stimindex graphs, each individual/peptide combination is a separate datapoint.

### Single-cell RNA sequencing

For single-cell RNA sequencing, PBMC of 6 donors with no history of SARS-CoV-2 infection and negative values for SARS-COV-2 IgG were stimulated with 1 µg/ml iCope peptide in the presence of purified anti-CD28 (clone CD28.2, BD Biosciences) and anti-CD40 (clone HB14, Miltenyi Biotec). CD4^+^ T cells were enriched by MACS (Miltenyi Biotec) and CD40L^+^4-1BB^+^ and CD40L^-^4-1BB^-^ cells FACS sorted using an Aria SORP (BD). The cells were loaded with a maximum concentration of 1000 cells/µl and a maximum cell number of 17,000 cells on a Chromium Chip G (10x Genomics). Gene expression and TCR libraries were generated according to the manufacturer’s instruction using the Chromium Next GEM single cell 5′Library and Gel bead Kit V1.1 and Chromium Single Cell V(D)J Enrichment Kit for human T cells (10x Genomics). Sequencing was conducted with a NovaSeq 6000 cartridge (Illumina) with 20,000 reads per cell for GEX libraries and 5,000 reads per cell for TCR libraries.

### Singe cell transcriptome analysis

Single cell RNA was mapped to reference genome GRCh38-2020-A and preprocessed using the Cell Ranger Software v6.1.2 (10x Genomics). Quality control and analysis of data was done in R 4.0.5 (R Core Team (2021). R: A language and environment for statistical computing. R Foundation for Statistical Computing, Vienna, Austria. URL https://www.R-project.org/) using the "Seurat" package (*73*). To remove low quality cells, doublets and empty cells thresholds were set to 840-3000 RNA features and less than 5% mitochondrial RNA. Data were normalized by using the LogNormalize function of the Seurat package and genes detected in less than 0.1% of the cells were excluded. For gene expression analysis the TCR genes were excluded from the data set to avoid TCR biased clustering. tSNE plots of CD40L^+^4-1BB^+^ or CD40L^-^4-1BB^-^ cells from young and older donors were overlaid using the in-built plot function of the Seurat package. Following gene signatures were analyzed: T naïve/memory (SELL, LEF1, TCD7, CCR7, CD27, CD28), cytotoxicity (PRF1, GNLY, NKG7, GZMB), cytokines (IL-2, IFN-γ, TNF-α, TGFβ1), senescence & exhaustion (LCK, ZAP70, LAT, TIGIT, CTLA4, BTLA, CD244, CD160, BATF, LCP2, TOX, PDCD1, KLRG1, HAVCR2 (TIM3), LAG3), anergy (EGR2, EGR3 DGK*α*, GRAIL, CD200) (*74–77*). Normalized and log2 transformed expression values of signature genes were shown in a heatmap using the DoHeatmap function of the Seurat package. Zero expression values were set to white color.

### Singe cell TCR analysis

Single cell TCR data were preprocessed using the Cell Ranger Software v6.1.2 (10x Genomics) and the GRCh38-2020-A reference genome. Data were further processed in R using the "immunarch" package (https://CRAN.R-project.org/package=immunarch, R package version 0.6.7.). Only cells which passed the quality controls in the gene expression analysis and containing exactly one TCR alpha and one TCR beta chain were used for further analysis. Diversity indices and rare clonal proportions were calculated using the corresponding functions of the immunarch package.

### Data analysis and statistics

Sequence data for analysis of the mutational landscape of SARS-CoV-2 were from the GISAID EpiCov database (https://www.epicov.org/). Study data were collected and managed using REDCap electronic data capture tools hosted at Charité (*78*, *79*). Flow cytometry data were analyzed with FlowJo 10.6 (FlowJo LLC) and statistical analysis conducted with GraphPad Prism 9. If not stated otherwise, data are plotted as mean. *N* indicates the number of donors. *P*-values were set as follows: \**P* <0.05, \*\**P* <0.01, and \*\*\**P*<0.001.

## Acknowledgements

The authors thank Zehra Uyar-Adin, Kübrah Gürcan, Luzie Bartsch, Maren Eckey, Pavlo Holenya and the CCC and CCP associates for their contributions to donor recruitment, sample processing and measurement. We would like to thank Desiree Kunkel, Jacqueline Keye and the BIH Flow & Mass Cytometry Core Facility for their support.

## Author contributions

Conceptualization: L.L. Investigation: L.L.

Data curation: L.L., K.J., U.R., L.H., J.B.

Visualization: L.L., K.J., U.R., J.K., F.K.

Formal Analysis: L.L., K.J., J.K., B.R., F.K.

Funding acquisition: A.T.

Resources: U.R., L.M-A., L.H., N.M., M.M., B.K., M.D., B.S., K.S., H.W., J.M., M.G., A.N., N.L., B.T., J.B., F.K.

Writing: L.L., C.G.-T., A.T.

**Competing interests**

The author U.R. was and F.K. is an employee, H.W. was the CEO of JPT. L.L., L.H., and A.T. are named on a filed patent application regarding the usage of CD3 downregulation as method for direct analysis of functional avidity of T cells and a patent application regarding the usage of iCope as method for the direct analysis of SARS-CoV-2 immune responses. All other authors declare that they have no competing interests.

**Data and materials availability**

All data are available in the main text or in the supplementary materials. single-cell RNA sequencing data is deposited under *TBA*.

**Supplementary Materials**

Figs. S1 to S5, Table S1 and S2

## References

1. COVID-19 Host Genetics Initiative, Mapping the human genetic architecture of COVID-19. Nature 600, 472–477 (2021).

2. A. B. Docherty, E. M. Harrison, C. A. Green, H. E. Hardwick, R. Pius, L. Norman, K. A. Holden, J. M. Read, F. Dondelinger, G. Carson, L. Merson, J. Lee, D. Plotkin, L. Sigfrid, S. Halpin, C. Jackson, C. Gamble, P. W. Horby, J. S. Nguyen-Van-Tam, A. Ho, C. D. Russell, J. Dunning, P. J. Openshaw, J. K. Baillie, M. G. Semple, Features of 20 133 UK patients in hospital with covid-19 using the ISARIC WHO Clinical Characterisation Protocol: prospective observational cohort study. BMJ 369, m1985 (2020).

3. F. Zhou, T. Yu, R. Du, G. Fan, Y. Liu, Z. Liu, J. Xiang, Y. Wang, B. Song, X. Gu, L. Guan, Y. Wei, H. Li, X. Wu, J. Xu, S. Tu, Y. Zhang, H. Chen, B. Cao, Clinical course and risk factors for mortality of adult inpatients with COVID-19 in Wuhan, China: a retrospective cohort study. The Lancet 395, 1054–1062 (2020).

4. R. Kundu, J. S. Narean, L. Wang, J. Fenn, T. Pillay, N. D. Fernandez, E. Conibear, A. Koycheva, M. Davies, M. Tolosa-Wright, S. Hakki, R. Varro, E. McDermott, S. Hammett, J. Cutajar, R. S. Thwaites, E. Parker, C. Rosadas, M. McClure, R. Tedder, G. P. Taylor, J. Dunning, A. Lalvani, Cross-reactive memory T cells associate with protection against SARS-CoV-2 infection in COVID-19 contacts. Nat Commun 13, 80 (2022).

5. L. Swadling, M. O. Diniz, N. M. Schmidt, O. E. Amin, A. Chandran, E. Shaw, C. Pade, J. M. Gibbons, N. Le Bert, A. T. Tan, A. Jeffery-Smith, C. C. S. Tan, C. Y. L. Tham, S. Kucykowicz, G. Aidoo-Micah, J. Rosenheim, J. Davies, M. Johnson, M. P. Jensen, G. Joy, L. E. McCoy, A. M. Valdes, B. M. Chain, D. Goldblatt, D. M. Altmann, R. J. Boyton, C. Manisty, T. A. Treibel, J. C. Moon, L. van Dorp, F. Balloux, Á. McKnight, M. Noursadeghi, A. Bertoletti, M. K. Maini, Pre-existing polymerase-specific T cells expand in abortive seronegative SARS-CoV-2. Nature, 1–10 (2021).

6. S. Seedat, H. Chemaitelly, H. H. Ayoub, M. Makhoul, G. R. Mumtaz, Z. Al Kanaani, A. Al Khal, E. Al Kuwari, A. A. Butt, P. Coyle, A. Jeremijenko, A. H. Kaleeckal, A. N. Latif, R. M. Shaik, H. M. Yassine, M. G. Al Kuwari, H. E. Al Romaihi, M. H. Al-Thani, R. Bertollini, L. J. Abu-Raddad, SARS-CoV-2 infection hospitalization, severity, criticality, and fatality rates in Qatar. Sci Rep 11, 18182 (2021).

7. A. J. Romero-Olmedo, A. R. Schulz, S. Hochstätter, D. Das Gupta, I. Virta, H. Hirseland, D. Staudenraus, B. Camara, C. Münch, V. Hefter, S. Sapre, V. Krähling, H. Müller-Kräuter, M. Widera, H. E. Mei, C. Keller, M. Lohoff, Induction of robust cellular and humoral immunity against SARS-CoV-2 after a third dose of BNT162b2 vaccine in previously unresponsive older adults. Nat Microbiol 7, 195–199 (2022).

8. L. Meyer-Arndt, T. Schwarz, L. Loyal, L. Henze, B. Kruse, M. Dingeldey, K. Gürcan, Z. Uyar-Aydin, M. A. Müller, C. Drosten, F. Paul, L. E. Sander, I. Demuth, R. Lauster, C. Giesecke-Thiel, J. Braun, V. M. Corman, A. Thiel, Cutting Edge: Serum but Not Mucosal Antibody Responses Are Associated with Pre-Existing SARS-CoV-2 Spike Cross-Reactive CD4+ T Cells following BNT162b2 Vaccination in the Elderly. The Journal of Immunology 208, 1001–1005 (2022).

9. N. Jo, Y. Hidaka, O. Kikuchi, M. Fukahori, T. Sawada, M. Aoki, M. Yamamoto, M. Nagao, S. Morita, T. E. Nakajima, M. Muto, Y. Hamazaki, Impaired CD4+ T cell response in older adults is associated with reduced immunogenicity and reactogenicity of mRNA COVID-19 vaccination. Nat Aging 3, 82–92 (2023).

10. S. Cele, L. Jackson, D. S. Khoury, K. Khan, T. Moyo-Gwete, H. Tegally, J. E. San, D. Cromer, C. Scheepers, D. G. Amoako, F. Karim, M. Bernstein, G. Lustig, D. Archary, M. Smith, Y. Ganga, Z. Jule, K. Reedoy, S.-H. Hwa, J. Giandhari, J. M. Blackburn, B. I. Gosnell, S. S. Abdool Karim, W. Hanekom, A. von Gottberg, J. N. Bhiman, R. J. Lessells, M.-Y. S. Moosa, M. P. Davenport, T. de Oliveira, P. L. Moore, A. Sigal, Omicron extensively but incompletely escapes Pfizer BNT162b2 neutralization. Nature 602, 654–656 (2022).

11. P. J. M. Brouwer, T. G. Caniels, K. van der Straten, J. L. Snitselaar, Y. Aldon, S. Bangaru, J. L. Torres, N. M. A. Okba, M. Claireaux, G. Kerster, A. E. H. Bentlage, M. M. van Haaren, D. Guerra, J. A. Burger, E. E. Schermer, K. D. Verheul, N. van der Velde, A. van der Kooi, J. van Schooten, M. J. van Breemen, T. P. L. Bijl, K. Sliepen, A. Aartse, R. Derking, I. Bontjer, N. A. Kootstra, W. J. Wiersinga, G. Vidarsson, B. L. Haagmans, A. B. Ward, G. J. de Bree, R. W. Sanders, M. J. van Gils, Potent neutralizing antibodies from COVID-19 patients define multiple targets of vulnerability. Science 369, 643–650 (2020).

12. H. Gruell, K. Vanshylla, M. Korenkov, P. Tober-Lau, M. Zehner, F. Münn, H. Janicki, M. Augustin, P. Schommers, L. E. Sander, F. Kurth, C. Kreer, F. Klein, SARS-CoV-2 Omicron sublineages exhibit distinct antibody escape patterns. Cell Host Microbe 30, 1231–1241.e6 (2022).

13. K. Sano, D. Bhavsar, G. Singh, D. Floda, K. Srivastava, C. Gleason, J. M. Carreño, V. Simon, F. Krammer, SARS-CoV-2 vaccination induces mucosal antibody responses in previously infected individuals. Nat Commun 13, 5135 (2022).

14. Y. Gao, C. Cai, A. Grifoni, T. R. Müller, J. Niessl, A. Olofsson, M. Humbert, L. Hansson, A. Österborg, P. Bergman, P. Chen, A. Olsson, J. K. Sandberg, D. Weiskopf, D. A. Price, H.-G. Ljunggren, A. C. Karlsson, A. Sette, S. Aleman, M. Buggert, Ancestral SARS-CoV-2-specific T cells cross-recognize the Omicron variant. Nat Med 28, 472–476 (2022).

15. L. Guo, G. Wang, Y. Wang, Q. Zhang, L. Ren, X. Gu, T. Huang, J. Zhong, Y. Wang, X. Wang, L. Huang, L. Xu, C. Wang, L. Chen, X. Xiao, Y. Peng, J. C. Knight, T. Dong, B. Cao, J. Wang, SARS-CoV-2-specific antibody and T-cell responses 1 year after infection in people recovered from COVID-19: a longitudinal cohort study. The Lancet Microbe 3, e348–e356 (2022).

16. A. Tarke, C. H. Coelho, Z. Zhang, J. M. Dan, E. D. Yu, N. Methot, N. I. Bloom, B. Goodwin, E. Phillips, S. Mallal, J. Sidney, G. Filaci, D. Weiskopf, R. da, S. Antunes, S. Crotty, A. Grifoni, A. Sette, SARS-CoV-2 vaccination induces immunological T cell memory able to cross-recognize variants from Alpha to Omicron. Cell 185, 847–859.e11 (2022).

17. A. Muik, B. G. Lui, J. Quandt, H. Diao, Y. Fu, M. Bacher, J. Gordon, A. Toker, J. Grosser, O. Ozhelvaci, K. Grikscheit, S. Hoehl, N. Kohmer, Y. Lustig, G. Regev-Yochay, S. Ciesek, K. Beguir, A. Poran, I. Vogler, Ö. Türeci, U. Sahin, Progressive loss of conserved spike protein neutralizing antibody sites in Omicron sublineages is balanced by preserved T cell immunity. Cell Reports 42, 112888 (2023).

18. L. Loyal, J. Braun, L. Henze, B. Kruse, M. Dingeldey, U. Reimer, F. Kern, T. Schwarz, M. Mangold, C. Unger, F. Dörfler, S. Kadler, J. Rosowski, K. Gürcan, Z. Uyar-Aydin, M. Frentsch, F. Kurth, K. Schnatbaum, M. Eckey, S. Hippenstiel, A. Hocke, M. A. Müller, B. Sawitzki, S. Miltenyi, F. Paul, M. A. Mall, H. Wenschuh, S. Voigt, C. Drosten, R. Lauster, N. Lachman, L.-E. Sander, V. M. Corman, J. Röhmel, L. Meyer-Arndt, A. Thiel, C. Giesecke-Thiel, Cross-reactive CD4+ T cells enhance SARS-CoV-2 immune responses upon infection and vaccination. Science 374, eabh1823 (2021).

19. J. Braun, L. Loyal, M. Frentsch, D. Wendisch, P. Georg, F. Kurth, S. Hippenstiel, M. Dingeldey, B. Kruse, F. Fauchere, E. Baysal, M. Mangold, L. Henze, R. Lauster, M. A. Mall, K. Beyer, J. Röhmel, S. Voigt, J. Schmitz, S. Miltenyi, I. Demuth, M. A. Müller, A. Hocke, M. Witzenrath, N. Suttorp, F. Kern, U. Reimer, H. Wenschuh, C. Drosten, V. M. Corman, C. Giesecke-Thiel, L. E. Sander, A. Thiel, SARS-CoV-2-reactive T cells in healthy donors and patients with COVID-19. Nature 587, 270–274 (2020).

20. S. M. Murray, A. M. Ansari, J. Frater, P. Klenerman, S. Dunachie, E. Barnes, A. Ogbe, The impact of pre-existing cross-reactive immunity on SARS-CoV-2 infection and vaccine responses. Nat Rev Immunol, 1–13 (2022).

21. J. Mateus, J. M. Dan, Z. Zhang, C. Rydyznski Moderbacher, M. Lammers, B. Goodwin, A. Sette, S. Crotty, D. Weiskopf, Low-dose mRNA-1273 COVID-19 vaccine generates durable memory enhanced by cross-reactive T cells. Science 0, eabj9853.

22. A. T. Tan, M. Linster, C. W. Tan, N. L. Bert, W. N. Chia, K. Kunasegaran, Y. Zhuang, C. Y. L. Tham, A. Chia, G. J. D. Smith, B. Young, S. Kalimuddin, J. G. H. Low, D. Lye, L.-F. Wang, A. Bertoletti, Early induction of functional SARS-CoV-2-specific T cells associates with rapid viral clearance and mild disease in COVID-19 patients. Cell Reports **34** (2021).

23. V. Mallajosyula, C. Ganjavi, S. Chakraborty, A. M. McSween, A. J. Pavlovitch-Bedzyk, J. Wilhelmy, A. Nau, M. Manohar, K. C. Nadeau, M. M. Davis, CD8+ T cells specific for conserved coronavirus epitopes correlate with milder disease in patients with COVID-19. Science Immunology 6, eabg5669 (2021).

24. M. M. Painter, T. S. Johnston, K. A. Lundgreen, J. J. S. Santos, J. S. Qin, R. R. Goel, S. A. Apostolidis, D. Mathew, B. Fulmer, J. C. Williams, M. L. McKeague, A. Pattekar, A. Goode, S. Nasta, A. E. Baxter, J. R. Giles, A. N. Skelly, L. E. Felley, M. McLaughlin, J. Weaver, Penn Medicine BioBank, O. Kuthuru, J. Dougherty, S. Adamski, S. Long, M. Kee, C. Clendenin, R. da Silva Antunes, A. Grifoni, D. Weiskopf, A. Sette, A. C. Huang, D. J. Rader, S. E. Hensley, P. Bates, A. R. Greenplate, E. J. Wherry, Prior vaccination promotes early activation of memory T cells and enhances immune responses during SARS-CoV-2 breakthrough infection. Nat Immunol 24, 1711–1724 (2023).

25. A. C. Dowell, M. S. Butler, E. Jinks, G. Tut, T. Lancaster, P. Sylla, J. Begum, R. Bruton, H. Pearce, K. Verma, N. Logan, G. Tyson, E. Spalkova, S. Margielewska-Davies, G. S. Taylor, E. Syrimi, F. Baawuah, J. Beckmann, I. O. Okike, S. Ahmad, J. Garstang, A. J. Brent, B. Brent, G. Ireland, F. Aiano, Z. Amin-Chowdhury, S. Jones, R. Borrow, E. Linley, J. Wright, R. Azad, D. Waiblinger, C. Davis, E. C. Thomson, M. Palmarini, B. J. Willett, W. S. Barclay, J. Poh, G. Amirthalingam, K. E. Brown, M. E. Ramsay, J. Zuo, P. Moss, S. Ladhani, Children develop robust and sustained cross-reactive spike-specific immune responses to SARS-CoV-2 infection. Nat Immunol 23, 40–49 (2022).

26. M. Humbert, A. Olofsson, D. Wullimann, J. Niessl, E. B. Hodcroft, C. Cai, Y. Gao, E. Sohlberg, R. Dyrdak, F. Mikaeloff, U. Neogi, J. Albert, K.-J. Malmberg, F. Lund-Johansen, S. Aleman, L. Björkhem-Bergman, M. C. Jenmalm, H.-G. Ljunggren, M. Buggert, A. C. Karlsson, Functional SARS-CoV-2 cross-reactive CD4+ T cells established in early childhood decline with age. Proceedings of the National Academy of Sciences 120, e2220320120 (2023).

27. F. Bianchini, V. Crivelli, M. E. Abernathy, C. Guerra, M. Palus, J. Muri, H. Marcotte, A. Piralla, M. Pedotti, R. De Gasparo, L. Simonelli, M. Matkovic, C. Toscano, M. Biggiogero, V. Calvaruso, P. Svoboda, T. Cervantes Rincón, T. Fava, L. Podešvová, A. A. Shanbhag, A. Celoria, J. Sgrignani, M. Stefanik, V. Hönig, V. Pranclova, T. Michalcikova, J. Prochazka, G. Guerrini, D. Mehn, A. Ciabattini, H. Abolhassani, D. Jarrossay, M. Uguccioni, D. Medaglini, Q. Pan-Hammarström, L. Calzolai, D. Fernandez, F. Baldanti, A. Franzetti-Pellanda, C. Garzoni, R. Sedlacek, D. Ruzek, L. Varani, A. Cavalli, C. O. Barnes, D. F. Robbiani, Human neutralizing antibodies to cold linear epitopes and subdomain 1 of the SARS-CoV-2 spike glycoprotein. Science Immunology 8, eade0958 (2023).

28. A. M. Johansson, U. Malhotra, Y. G. Kim, R. Gomez, M. P. Krist, A. Wald, D. M. Koelle, W. W. Kwok, Cross-reactive and mono-reactive SARS-CoV-2 CD4+ T cells in prepandemic and COVID-19 convalescent individuals. PLOS Pathogens 17, e1010203 (2021).

29. J. S. Low, D. Vaqueirinho, F. Mele, M. Foglierini, J. Jerak, M. Perotti, D. Jarrossay, S. Jovic, L. Perez, R. Cacciatore, T. Terrot, A. F. Pellanda, M. Biggiogero, C. Garzoni, P. Ferrari, A. Ceschi, A. Lanzavecchia, F. Sallusto, A. Cassotta, Clonal analysis of immunodominance and cross-reactivity of the CD4 T cell response to SARS-CoV-2. Science 372, 1336–1341 (2021).

30. E. Shrock, E. Fujimura, T. Kula, R. T. Timms, I.-H. Lee, Y. Leng, M. L. Robinson, B. M. Sie, M. Z. Li, Y. Chen, J. Logue, A. Zuiani, D. McCulloch, F. J. N. Lelis, S. Henson, D. R. Monaco, M. Travers, S. Habibi, W. A. Clarke, P. Caturegli, O. Laeyendecker, A. Piechocka-Trocha, J. Z. Li, A. Khatri, H. Y. Chu, M. C.-19 Collection &, P. Team16‡, A.-C. Villani, K. Kays, M. B. Goldberg, N. Hacohen, M. R. Filbin, X. G. Yu, B. D. Walker, D. R. Wesemann, H. B. Larman, J. A. Lederer, S. J. Elledge, Viral epitope profiling of COVID-19 patients reveals cross-reactivity and correlates of severity. Science **370** (2020).

31. J. T. Ladner, S. N. Henson, A. S. Boyle, A. L. Engelbrektson, Z. W. Fink, F. Rahee, J. D’ambrozio, K. E. Schaecher, M. Stone, W. Dong, S. Dadwal, J. Yu, M. A. Caligiuri, P. Cieplak, M. Bjørås, M. H. Fenstad, S. A. Nordbø, D. E. Kainov, N. Muranaka, M. S. Chee, S. A. Shiryaev, J. A. Altin, Epitope-resolved profiling of the SARS-CoV-2 antibody response identifies cross-reactivity with endemic human coronaviruses. Cell Reports Medicine 2, 100189 (2021).

32. L. Guo, S. Lin, Z. Chen, Y. Cao, B. He, G. Lu, Targetable elements in SARS-CoV-2 S2 subunit for the design of pan-coronavirus fusion inhibitors and vaccines. Sig Transduct Target Ther 8, 1–22 (2023).

33. Z. Wang, F. Muecksch, D. Schaefer-Babajew, S. Finkin, C. Viant, C. Gaebler, H.-H. Hoffmann, C. O. Barnes, M. Cipolla, V. Ramos, T. Y. Oliveira, A. Cho, F. Schmidt, J. Da Silva, E. Bednarski, L. Aguado, J. Yee, M. Daga, M. Turroja, K. G. Millard, M. Jankovic, A. Gazumyan, Z. Zhao, C. M. Rice, P. D. Bieniasz, M. Caskey, T. Hatziioannou, M. C. Nussenzweig, Naturally enhanced neutralizing breadth against SARS-CoV-2 one year after infection. Nature 595, 426–431 (2021).

34. Y. Cao, J. Wang, F. Jian, T. Xiao, W. Song, A. Yisimayi, W. Huang, Q. Li, P. Wang, R. An, J. Wang, Y. Wang, X. Niu, S. Yang, H. Liang, H. Sun, T. Li, Y. Yu, Q. Cui, S. Liu, X. Yang, S. Du, Z. Zhang, X. Hao, F. Shao, R. Jin, X. Wang, J. Xiao, Y. Wang, X. S. Xie, Omicron escapes the majority of existing SARS-CoV-2 neutralizing antibodies. Nature 602, 657–663 (2022).

35. H. Karsten, L. Cords, T. Westphal, M. Knapp, T. T. Brehm, L. Hermanussen, T. F. Omansen, S. Schmiedel, R. Woost, V. Ditt, S. Peine, M. Lütgehetmann, S. Huber, C. Ackermann, M. Wittner, M. M. Addo, A. Sette, J. Sidney, J. Schulze zur Wiesch, High-resolution analysis of individual spike peptide-specific CD4+ T-cell responses in vaccine recipients and COVID-19 patients. Clinical & Translational Immunology 11, e1410 (2022).

36. C. J. Reynolds, C. Pade, J. M. Gibbons, A. D. Otter, K.-M. Lin, D. Muñoz Sandoval, F. P. Pieper, D. K. Butler, S. Liu, G. Joy, N. Forooghi, T. A. Treibel, C. Manisty, J. C. Moon, COVIDsortium Investigators, COVIDsortium Immune Correlates Network, A. Semper, T. Brooks, Á. McKnight, D. M. Altmann, R. J. Boyton, Immune boosting by B.1.1.529 (Omicron) depends on previous SARS-CoV-2 exposure. Science **377**, eabq1841 (2022).

37. E. J. Yager, M. Ahmed, K. Lanzer, T. D. Randall, D. L. Woodland, M. A. Blackman, Age-associated decline in T cell repertoire diversity leads to holes in the repertoire and impaired immunity to influenza virus. Journal of Experimental Medicine 205, 711–723 (2008).

38. B. V. Kumar, T. J. Connors, D. L. Farber, Human T Cell Development, Localization, and Function throughout Life. Immunity 48, 202–213 (2018).

39. X. Sun, T. Nguyen, A. Achour, A. Ko, J. Cifello, C. Ling, J. Sharma, T. Hiroi, Y. Zhang, C. W. Chia, W. W. III, W. W. Wu, L. Zukley, J.-N. Phue, K. G. Becker, R.-F. Shen, L. Ferrucci, N. Weng, Longitudinal analysis reveals age-related changes in the T cell receptor repertoire of human T cell subsets (2022). 10.1172/JCI158122.

40. Q. Qi, Y. Liu, Y. Cheng, J. Glanville, D. Zhang, J.-Y. Lee, R. A. Olshen, C. M. Weyand, S. D. Boyd, J. J. Goronzy, Diversity and clonal selection in the human T-cell repertoire. PNAS 111, 13139–13144 (2014).

41. A. R. Schulz, J. N. Mälzer, C. Domingo, K. Jürchott, A. Grützkau, N. Babel, M. Nienen, T. Jelinek, M. Niedrig, A. Thiel, Low Thymic Activity and Dendritic Cell Numbers Are Associated with the Immune Response to Primary Viral Infection in Elderly Humans. The Journal of Immunology 195, 4699–4711 (2015).

42. O. V. Britanova, M. Shugay, E. M. Merzlyak, D. B. Staroverov, E. V. Putintseva, M. A. Turchaninova, I. Z. Mamedov, M. V. Pogorelyy, D. A. Bolotin, M. Izraelson, A. N. Davydov, E. S. Egorov, S. A. Kasatskaya, D. V. Rebrikov, S. Lukyanov, D. M. Chudakov, Dynamics of Individual T Cell Repertoires: From Cord Blood to Centenarians. J Immunol, 1600005 (2016).

43. A. Nelde, T. Bilich, J. S. Heitmann, Y. Maringer, H. R. Salih, M. Roerden, M. Lübke, J. Bauer, J. Rieth, M. Wacker, A. Peter, S. Hörber, B. Traenkle, P. D. Kaiser, U. Rothbauer, M. Becker, D. Junker, G. Krause, M. Strengert, N. Schneiderhan-Marra, M. F. Templin, T. O. Joos, D. J. Kowalewski, V. Stos-Zweifel, M. Fehr, A. Rabsteyn, V. Mirakaj, J. Karbach, E. Jäger, M. Graf, L.-C. Gruber, D. Rachfalski, B. Preuß, I. Hagelstein, M. Märklin, T. Bakchoul, C. Gouttefangeas, O. Kohlbacher, R. Klein, S. Stevanović, H.-G. Rammensee, J. S. Walz, SARS-CoV-2-derived peptides define heterologous and COVID-19-induced T cell recognition. Nature Immunology 22, 74–85 (2021).

44. Y. Peng, A. J. Mentzer, G. Liu, X. Yao, Z. Yin, D. Dong, W. Dejnirattisai, T. Rostron, P. Supasa, C. Liu, C. López-Camacho, J. Slon-Campos, Y. Zhao, D. I. Stuart, G. C. Paesen, J. M. Grimes, A. A. Antson, O. W. Bayfield, D. E. D. P. Hawkins, D.-S. Ker, B. Wang, L. Turtle, K. Subramaniam, P. Thomson, P. Zhang, C. Dold, J. Ratcliff, P. Simmonds, T. de Silva, P. Sopp, D. Wellington, U. Rajapaksa, Y.-L. Chen, M. Salio, G. Napolitani, W. Paes, P. Borrow, B. M. Kessler, J. W. Fry, N. F. Schwabe, M. G. Semple, J. K. Baillie, S. C. Moore, P. J. M. Openshaw, M. A. Ansari, S. Dunachie, E. Barnes, J. Frater, G. Kerr, P. Goulder, T. Lockett, R. Levin, Y. Zhang, R. Jing, L.-P. Ho, R. J. Cornall, C. P. Conlon, P. Klenerman, G. R. Screaton, J. Mongkolsapaya, A. McMichael, J. C. Knight, G. Ogg, T. Dong, Broad and strong memory CD4+ and CD8+ T cells induced by SARS-CoV-2 in UK convalescent individuals following COVID-19. Nat Immunol 21, 1336–1345 (2020).

45. D. A. Collier, I. A. T. M. Ferreira, P. Kotagiri, R. P. Datir, E. Y. Lim, E. Touizer, B. Meng, A. Abdullahi, A. Elmer, N. Kingston, B. Graves, E. Le Gresley, D. Caputo, L. Bergamaschi, K. G. C. Smith, J. R. Bradley, L. Ceron-Gutierrez, P. Cortes-Acevedo, G. Barcenas-Morales, M. A. Linterman, L. E. McCoy, C. Davis, E. Thomson, P. A. Lyons, E. McKinney, R. Doffinger, M. Wills, R. K. Gupta, Age-related immune response heterogeneity to SARS-CoV-2 vaccine BNT162b2. Nature 596, 417–422 (2021).

46. T. A. Schwickert, B. Alabyev, T. Manser, M. C. Nussenzweig, Germinal center reutilization by newly activated B cells. Journal of Experimental Medicine 206, 2907–2914 (2009).

47. T. A. Schwickert, G. D. Victora, D. R. Fooksman, A. O. Kamphorst, M. R. Mugnier, A. D. Gitlin, M. L. Dustin, M. C. Nussenzweig, A dynamic T cell–limited checkpoint regulates affinity-dependent B cell entry into the germinal center. Journal of Experimental Medicine 208, 1243–1252 (2011).

48. K. W. Ng, N. Faulkner, G. H. Cornish, A. Rosa, R. Harvey, S. Hussain, R. Ulferts, C. Earl, A. G. Wrobel, D. J. Benton, C. Roustan, W. Bolland, R. Thompson, A. Agua-Doce, P. Hobson, J. Heaney, H. Rickman, S. Paraskevopoulou, C. F. Houlihan, K. Thomson, E. Sanchez, G. Y. Shin, M. J. Spyer, D. Joshi, N. O’Reilly, P. A. Walker, S. Kjaer, A. Riddell, C. Moore, B. R. Jebson, M. Wilkinson, L. R. Marshall, E. C. Rosser, A. Radziszewska, H. Peckham, C. Ciurtin, L. R. Wedderburn, R. Beale, C. Swanton, S. Gandhi, B. Stockinger, J. McCauley, S. J. Gamblin, L. E. McCoy, P. Cherepanov, E. Nastouli, G. Kassiotis, Preexisting and de novo humoral immunity to SARS-CoV-2 in humans. Science 370, 1339–1343 (2020).

49. G. Song, W.-T. He, S. Callaghan, F. Anzanello, D. Huang, J. Ricketts, J. L. Torres, N. Beutler, L. Peng, S. Vargas, J. Cassell, M. Parren, L. Yang, C. Ignacio, D. M. Smith, J. E. Voss, D. Nemazee, A. B. Ward, T. Rogers, D. R. Burton, R. Andrabi, Cross-reactive serum and memory B-cell responses to spike protein in SARS-CoV-2 and endemic coronavirus infection. Nat Commun 12, 2938 (2021).

50. R. C. Brewer, N. S. Ramadoss, L. J. Lahey, S. Jahanbani, W. H. Robinson, T. V. Lanz, BNT162b2 vaccine induces divergent B cell responses to SARS-CoV-2 S1 and S2. Nat Immunol 23, 33–39 (2022).

51. K. W. Ng, N. Faulkner, K. Finsterbusch, M. Wu, R. Harvey, S. Hussain, M. Greco, Y. Liu, S. Kjaer, C. Swanton, S. Gandhi, R. Beale, S. J. Gamblin, P. Cherepanov, J. McCauley, R. Daniels, M. Howell, H. Arase, A. Wack, D. L. V. Bauer, G. Kassiotis, SARS-CoV-2 S2–targeted vaccination elicits broadly neutralizing antibodies. Science Translational Medicine 14, eabn3715 (2022).

52. C. Dacon, C. Tucker, L. Peng, C.-C. D. Lee, T.-H. Lin, M. Yuan, Y. Cong, L. Wang, L. Purser, J. K. Williams, C.-W. Pyo, I. Kosik, Z. Hu, M. Zhao, D. Mohan, A. J. R. Cooper, M. Peterson, J. Skinner, S. Dixit, E. Kollins, L. Huzella, D. Perry, R. Byrum, S. Lembirik, D. Drawbaugh, B. Eaton, Y. Zhang, E. S. Yang, M. Chen, K. Leung, R. S. Weinberg, A. Pegu, D. E. Geraghty, E. Davidson, I. Douagi, S. Moir, J. W. Yewdell, C. Schmaljohn, P. D. Crompton, M. R. Holbrook, D. Nemazee, J. R. Mascola, I. A. Wilson, J. Tan, Broadly neutralizing antibodies target the coronavirus fusion peptide. Science 377, 728–735 (2022).

53. J. S. Low, J. Jerak, M. A. Tortorici, M. McCallum, D. Pinto, A. Cassotta, M. Foglierini, F. Mele, R. Abdelnabi, B. Weynand, J. Noack, M. Montiel-Ruiz, S. Bianchi, F. Benigni, N. Sprugasci, A. Joshi, J. E. Bowen, C. Stewart, M. Rexhepaj, A. C. Walls, D. Jarrossay, D. Morone, P. Paparoditis, C. Garzoni, P. Ferrari, A. Ceschi, J. Neyts, L. A. Purcell, G. Snell, D. Corti, A. Lanzavecchia, D. Veesler, F. Sallusto, ACE2-binding exposes the SARS-CoV-2 fusion peptide to broadly neutralizing coronavirus antibodies. Science 377, 735–742 (2022).

54. D. Pinto, M. M. Sauer, N. Czudnochowski, J. S. Low, M. A. Tortorici, M. P. Housley, J. Noack, A. C. Walls, J. E. Bowen, B. Guarino, L. E. Rosen, J. di Iulio, J. Jerak, H. Kaiser, S. Islam, S. Jaconi, N. Sprugasci, K. Culap, R. Abdelnabi, C. Foo, L. Coelmont, I. Bartha, S. Bianchi, C. Silacci-Fregni, J. Bassi, R. Marzi, E. Vetti, A. Cassotta, A. Ceschi, P. Ferrari, P. E. Cippà, O. Giannini, S. Ceruti, C. Garzoni, A. Riva, F. Benigni, E. Cameroni, L. Piccoli, M. S. Pizzuto, M. Smithey, D. Hong, A. Telenti, F. A. Lempp, J. Neyts, C. Havenar-Daughton, A. Lanzavecchia, F. Sallusto, G. Snell, H. W. Virgin, M. Beltramello, D. Corti, D. Veesler, Broad betacoronavirus neutralization by a stem helix–specific human antibody. Science 373, 1109–1116 (2021).

55. P. Zhou, G. Song, H. Liu, M. Yuan, W. He, N. Beutler, X. Zhu, L. V. Tse, D. R. Martinez, A. Schäfer, F. Anzanello, P. Yong, L. Peng, K. Dueker, R. Musharrafieh, S. Callaghan, T. Capozzola, O. Limbo, M. Parren, E. Garcia, S. A. Rawlings, D. M. Smith, D. Nemazee, J. G. Jardine, Y. Safonova, B. Briney, T. F. Rogers, I. A. Wilson, R. S. Baric, L. E. Gralinski, D. R. Burton, R. Andrabi, Broadly neutralizing anti-S2 antibodies protect against all three human betacoronaviruses that cause deadly disease. Immunity 56, 669–686.e7 (2023).

56. T. M. Wilkinson, C. K. F. Li, C. S. C. Chui, A. K. Y. Huang, M. Perkins, J. C. Liebner, R. Lambkin-Williams, A. Gilbert, J. Oxford, B. Nicholas, K. J. Staples, T. Dong, D. C. Douek, A. J. McMichael, X.-N. Xu, Preexisting influenza-specific CD4+ T cells correlate with disease protection against influenza challenge in humans. Nat Med 18, 274–280 (2012).

57. S. Sridhar, S. Begom, A. Bermingham, K. Hoschler, W. Adamson, W. Carman, T. Bean, W. Barclay, J. J. Deeks, A. Lalvani, Cellular immune correlates of protection against symptomatic pandemic influenza. Nat Med 19, 1305–1312 (2013).

58. M. Nienen, U. Stervbo, F. Mölder, S. Kaliszczyk, L. Kuchenbecker, L. Gayova, B. Schweiger, K. Jürchott, J. Hecht, A. U. Neumann, S. Rahmann, T. Westhoff, P. Reinke, A. Thiel, N. Babel, The Role of Pre-existing Cross-Reactive Central Memory CD4 T-Cells in Vaccination With Previously Unseen Influenza Strains. Front. Immunol. 0 (2019).

59. K. G. Lanzer, L. L. Johnson, D. L. Woodland, M. A. Blackman, Impact of ageing on the response and repertoire of influenza virus-specific CD4 T cells. Immunity & Ageing 11, 9 (2014).

60. C. E. van de Sandt, T. H. O. Nguyen, N. A. Gherardin, J. C. Crawford, J. Samir, A. A. Minervina, M. V. Pogorelyy, S. Rizzetto, C. Szeto, J. Kaur, N. Ranson, S. Sonda, A. Harper, S. J. Redmond, H. A. McQuilten, T. Menon, S. Sant, X. Jia, K. Pedrina, T. Karapanagiotidis, N. Cain, S. Nicholson, Z. Chen, R. Lim, E. B. Clemens, A. Eltahla, N. L. La Gruta, J. Crowe, M. Lappas, J. Rossjohn, D. I. Godfrey, P. G. Thomas, S. Gras, K. L. Flanagan, F. Luciani, K. Kedzierska, Newborn and child-like molecular signatures in older adults stem from TCR shifts across human lifespan. Nat Immunol 24, 1890–1907 (2023).

61. M. Á. Palacios-Pedrero, J. M. Jansen, C. Blume, N. Stanislawski, R. Jonczyk, A. Molle, M. G. Hernandez, F. K. Kaiser, K. Jung, A. D. M. E. Osterhaus, G. F. Rimmelzwaan, G. Saletti, Signs of immunosenescence correlate with poor outcome of mRNA COVID-19 vaccination in older adults. Nat Aging 2, 896–905 (2022).

62. L. Loyal, S. Warth, K. Jürchott, F. Mölder, C. Nikolaou, N. Babel, M. Nienen, S. Durlanik, R. Stark, B. Kruse, M. Frentsch, R. Sabat, K. Wolk, A. Thiel, SLAMF7 and IL-6R define distinct cytotoxic versus helper memory CD8+ T cells. Nat Commun 11, 6357 (2020).

63. K. Hashimoto, T. Kouno, T. Ikawa, N. Hayatsu, Y. Miyajima, H. Yabukami, T. Terooatea, T. Sasaki, T. Suzuki, M. Valentine, G. Pascarella, Y. Okazaki, H. Suzuki, J. W. Shin, A. Minoda, I. Taniuchi, H. Okano, Y. Arai, N. Hirose, P. Carninci, Single-cell transcriptomics reveals expansion of cytotoxic CD4 T cells in supercentenarians. Proceedings of the National Academy of Sciences 116, 24242–24251 (2019).

64. M. Yoshida, K. B. Worlock, N. Huang, R. G. H. Lindeboom, C. R. Butler, N. Kumasaka, C. Dominguez Conde, L. Mamanova, L. Bolt, L. Richardson, K. Polanski, E. Madissoon, J. L. Barnes, J. Allen-Hyttinen, E. Kilich, B. C. Jones, A. de Wilton, A. Wilbrey-Clark, W. Sungnak, J. P. Pett, J. Weller, E. Prigmore, H. Yung, P. Mehta, A. Saleh, A. Saigal, V. Chu, J. M. Cohen, C. Cane, A. Iordanidou, S. Shibuya, A.-K. Reuschl, I. T. Herczeg, A. C. Argento, R. G. Wunderink, S. B. Smith, T. A. Poor, C. A. Gao, J. E. Dematte, G. Reynolds, M. Haniffa, G. S. Bowyer, M. Coates, M. R. Clatworthy, F. J. Calero-Nieto, B. Göttgens, C. O’Callaghan, N. J. Sebire, C. Jolly, P. De Coppi, C. M. Smith, A. V. Misharin, S. M. Janes, S. A. Teichmann, M. Z. Nikolić, K. B. Meyer, Local and systemic responses to SARS-CoV-2 infection in children and adults. Nature 602, 321–327 (2022).

65. B. J. Meckiff, C. Ramírez-Suástegui, V. Fajardo, S. J. Chee, A. Kusnadi, H. Simon, S. Eschweiler, A. Grifoni, E. Pelosi, D. Weiskopf, A. Sette, F. Ay, G. Seumois, C. H. Ottensmeier, P. Vijayanand, Imbalance of Regulatory and Cytotoxic SARS-CoV-2-Reactive CD4+ T Cells in COVID-19. Cell 183, 1340–1353.e16 (2020).

66. N. van Doremalen, J. N. Purushotham, J. E. Schulz, M. G. Holbrook, T. Bushmaker, A. Carmody, J. R. Port, C. K. Yinda, A. Okumura, G. Saturday, F. Amanat, F. Krammer, P. W. Hanley, B. J. Smith, J. Lovaglio, S. L. Anzick, K. Barbian, C. Martens, S. C. Gilbert, T. Lambe, V. J. Munster, Intranasal ChAdOx1 nCoV-19/AZD1222 vaccination reduces viral shedding after SARS-CoV-2 D614G challenge in preclinical models. Science Translational Medicine 13, eabh0755 (2021).

67. J. M. Carreño, A. Raskin, G. Singh, J. Tcheou, H. Kawabata, C. Gleason, K. Srivastava, V. Vigdorovich, N. Dambrauskas, S. L. Gupta, I. Gonzalez, J. L. Martinez, S. Slamanig, D. N. Sather, R. Raghunandan, P. Wirachwong, S. Muangnoicharoen, P. Pitisuttithum, J. Wrammert, M. S. Suthar, W. Sun, P. Palese, A. García-Sastre, V. Simon, F. Krammer, “The inactivated NDV-HXP-S COVID-19 vaccine induces a significantly higher ratio of neutralizing to non-neutralizing antibodies in humans as compared to mRNA vaccines” (preprint, Infectious Diseases (except HIV/AIDS), 2022); 10.1101/2022.01.25.22269808.

68. M. Madhavan, A. J. Ritchie, J. Aboagye, D. Jenkin, S. Provstgaad-Morys, I. Tarbet, D. Woods, S. Davies, M. Baker, A. Platt, A. Flaxman, H. Smith, S. Belij-Rammerstorfer, D. Wilkins, E. J. Kelly, T. Villafana, J. A. Green, I. Poulton, T. Lambe, A. V. S. Hill, K. J. Ewer, A. D. Douglas, Tolerability and immunogenicity of an intranasally-administered adenovirus-vectored COVID-19 vaccine: An open-label partially-randomised ascending dose phase I trial. eBioMedicine 0 (2022).

69. J. S. Heitmann, T. Bilich, C. Tandler, A. Nelde, Y. Maringer, M. Marconato, J. Reusch, S. Jäger, M. Denk, M. Richter, L. Anton, L. M. Weber, M. Roerden, J. Bauer, J. Rieth, M. Wacker, S. Hörber, A. Peter, C. Meisner, I. Fischer, M. W. Löffler, J. Karbach, E. Jäger, R. Klein, H.-G. Rammensee, H. R. Salih, J. S. Walz, A COVID-19 peptide vaccine for the induction of SARS-CoV-2 T cell immunity. Nature 601, 617–622 (2022).

70. M. O. Diniz, E. Mitsi, L. Swadling, J. Rylance, M. Johnson, D. Goldblatt, D. Ferreira, M. K. Maini, Airway-resident T cells from unexposed individuals cross-recognize SARS-CoV-2. Nat Immunol 23, 1324–1329 (2022).

71. J. Niessl, T. Sekine, J. Lange, V. Konya, M. Forkel, J. Maric, A. Rao, L. Mazzurana, E. Kokkinou, W. Weigel, S. Llewellyn-Lacey, E. B. Hodcroft, A. C. Karlsson, J. Fehrm, J. Sundman, D. A. Price, J. Mjösberg, D. Friberg, M. Buggert, Identification of resident memory CD8+ T cells with functional specificity for SARS-CoV-2 in unexposed oropharyngeal lymphoid tissue. Science Immunology 0, eabk0894.

72. J. Michel, M. Neumann, E. Krause, T. Rinner, T. Muzeniek, M. Grossegesse, G. Hille, F. Schwarz, A. Puyskens, S. Förster, B. Biere, D. Bourquain, C. Domingo, A. Brinkmann, L. Schaade, L. Schrick, A. Nitsche, Resource-efficient internally controlled in-house real-time PCR detection of SARS-CoV-2. Virology Journal 18, 110 (2021).

73. Y. Hao, S. Hao, E. Andersen-Nissen, W. M. Mauck, S. Zheng, A. Butler, M. J. Lee, A. J. Wilk, C. Darby, M. Zager, P. Hoffman, M. Stoeckius, E. Papalexi, E. P. Mimitou, J. Jain, A. Srivastava, T. Stuart, L. M. Fleming, B. Yeung, A. J. Rogers, J. M. McElrath, C. A. Blish, R. Gottardo, P. Smibert, R. Satija, Integrated analysis of multimodal single-cell data. Cell 184, 3573–3587.e29 (2021).

74. M. Safford, S. Collins, M. A. Lutz, A. Allen, C.-T. Huang, J. Kowalski, A. Blackford, M. R. Horton, C. Drake, R. H. Schwartz, J. D. Powell, Egr-2 and Egr-3 are negative regulators of T cell activation. Nat Immunol 6, 472–480 (2005).

75. A. Trefzer, P. Kadam, S.-H. Wang, S. Pennavaria, B. Lober, B. Akçabozan, J. Kranich, T. Brocker, N. Nakano, M. Irmler, J. Beckers, T. Straub, R. Obst, Dynamic adoption of anergy by antigen-exhausted CD4+ T cells. Cell Reports 34, 108748 (2021).

76. J.-Y. Zhang, X.-M. Wang, X. Xing, Z. Xu, C. Zhang, J.-W. Song, X. Fan, P. Xia, J.-L. Fu, S.-Y. Wang, R.- N. Xu, X.-P. Dai, L. Shi, L. Huang, T.-J. Jiang, M. Shi, Y. Zhang, A. Zumla, M. Maeurer, F. Bai, F.-S. Wang, Single-cell landscape of immunological responses in patients with COVID-19. Nat Immunol 21, 1107–1118 (2020).

77. Y. Zhao, Q. Shao, G. Peng, Exhaustion and senescence: two crucial dysfunctional states of T cells in the tumor microenvironment. Cell Mol Immunol 17, 27–35 (2020).

78. P. A. Harris, R. Taylor, R. Thielke, J. Payne, N. Gonzalez, J. G. Conde, Research electronic data capture (REDCap)—A metadata-driven methodology and workflow process for providing translational research informatics support. Journal of Biomedical Informatics 42, 377–381 (2009).

79. P. A. Harris, R. Taylor, B. L. Minor, V. Elliott, M. Fernandez, L. O’Neal, L. McLeod, G. Delacqua, F. Delacqua, J. Kirby, S. N. Duda, The REDCap consortium: Building an international community of software platform partners. Journal of Biomedical Informatics 95, 103208 (2019).

