## Supplementary Materials for "Aging and viral evolution impair immunity against dominant pan-coronavirus-reactive T cell epitope"

**Supplementary Materials:** Figs. S1 to S5, Table S1 and S2

### Figure S1

A

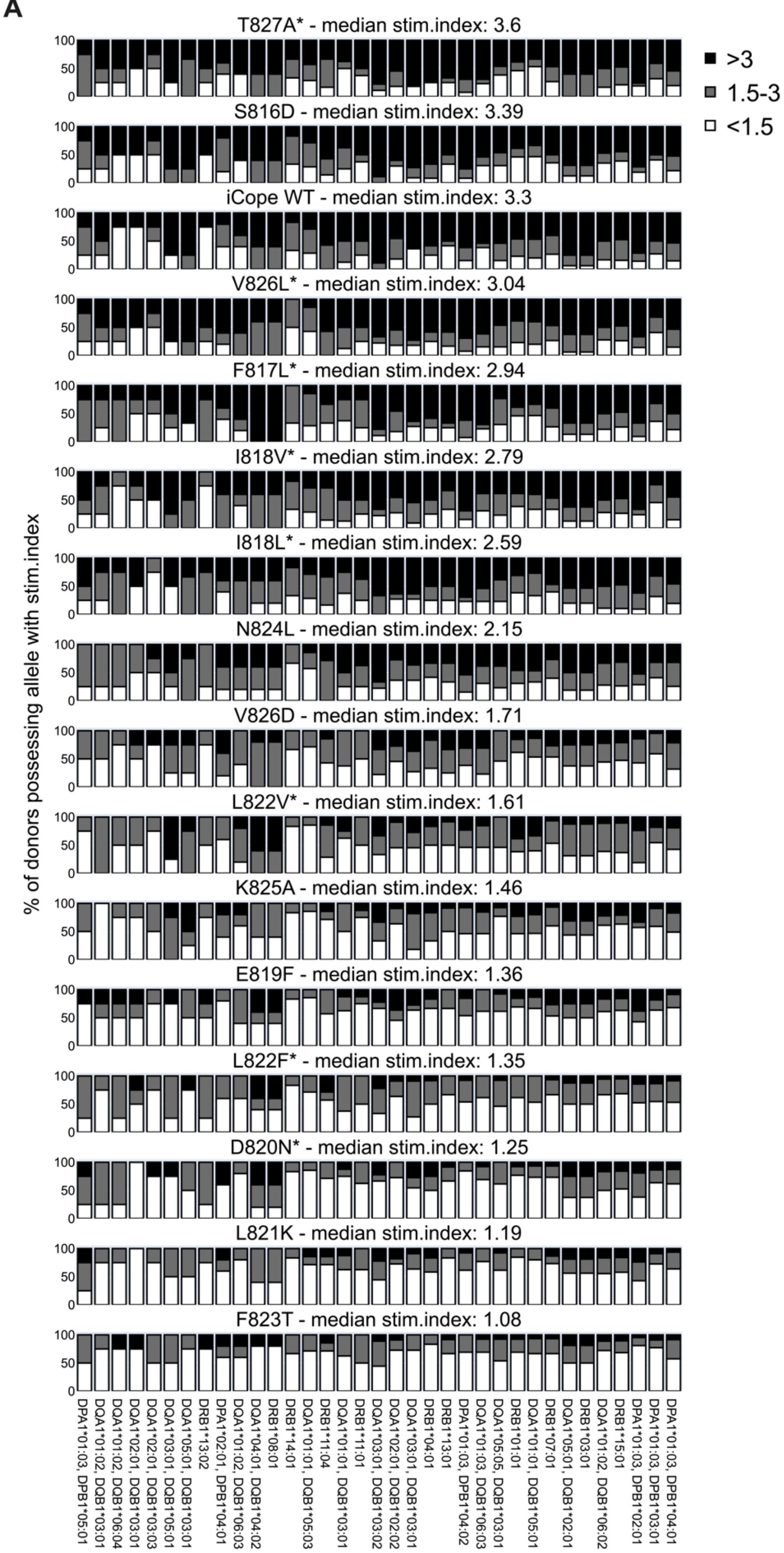
T827A* - median stim.index: 3.6
S816D - median stim.index: 3.39
iCope WT - median stim.index: 3.3
V826L* - median stim.index: 3.04
F817L* - median stim.index: 2.94
I818V* - median stim.index: 2.79
I818L* - median stim.index: 2.59
N824L - median stim.index: 2.15
V826D - median stim.index: 1.71
L822V* - median stim.index: 1.61
K825A - median stim.index: 1.46
E819F - median stim.index: 1.36
L822F* - median stim.index: 1.35
D820N* - median stim.index: 1.25
L821K - median stim.index: 1.19
F823T - median stim.index: 1.08
>3
1.5-3
<1.5
% of donors possessing allele with stim.index
DPA1*01:03, DPB1*05:01
DQA1*01:02, DQB1*03:01
DQA1*01:02, DQB1*06:04
DQA1*02:01, DQB1*03:01
DQA1*02:01, DQB1*03:03
DQA1*03:01, DQB1*05:01
DQA1*05:01, DQB1*03:01
DRB1*13:02
DPA1*02:01, DPB1*04:01
DQA1*01:02, DQB1*06:03
DQA1*04:01, DQB1*04:02
DRB1*08:01
DRB1*14:01
DQA1*01:01, DQB1*05:03
DRB1*11:04
DQA1*01:01, DQB1*03:01
DRB1*11:01
DQA1*03:01, DQB1*03:02
DQA1*02:01, DQB1*02:02
DQA1*03:01, DQB1*03:01
DRB1*04:01
DRB1*13:01
DPA1*01:03, DPB1*04:02
DQA1*01:03, DQB1*06:03
DQA1*05:05, DQB1*03:01
DRB1*01:01
DQA1*01:01, DQB1*05:01
DRB1*07:01
DQA1*05:01, DQB1*02:01
DRB1*03:01
DQA1*01:02, DQB1*06:02
DRB1*15:01
DPA1*01:03, DPB1*02:01
DPA1*01:03, DPB1*03:01
DPA1*01:03, DPB1*04:01

**Fig. S1: Impact of different alleles on T cell responsiveness against iCope mutations.** Lines indicate the percentage of different stim responses (y-axis) for all donors who possess the given allele (x-axis). Alleles with frequencies > 5% in the analyzed cohort (all donors of Table 1) are shown. Each donor is represented with at least one allele combination. The color code indicates the percentage of donors responding to the T cell stimulation with the respective peptide with a stimulation index < 1.5 (white, unresponsive), 1.5-3 (grey, weak response) or > 3 (black, strong response). * indicates documented iCope mutations.

### Figure S2

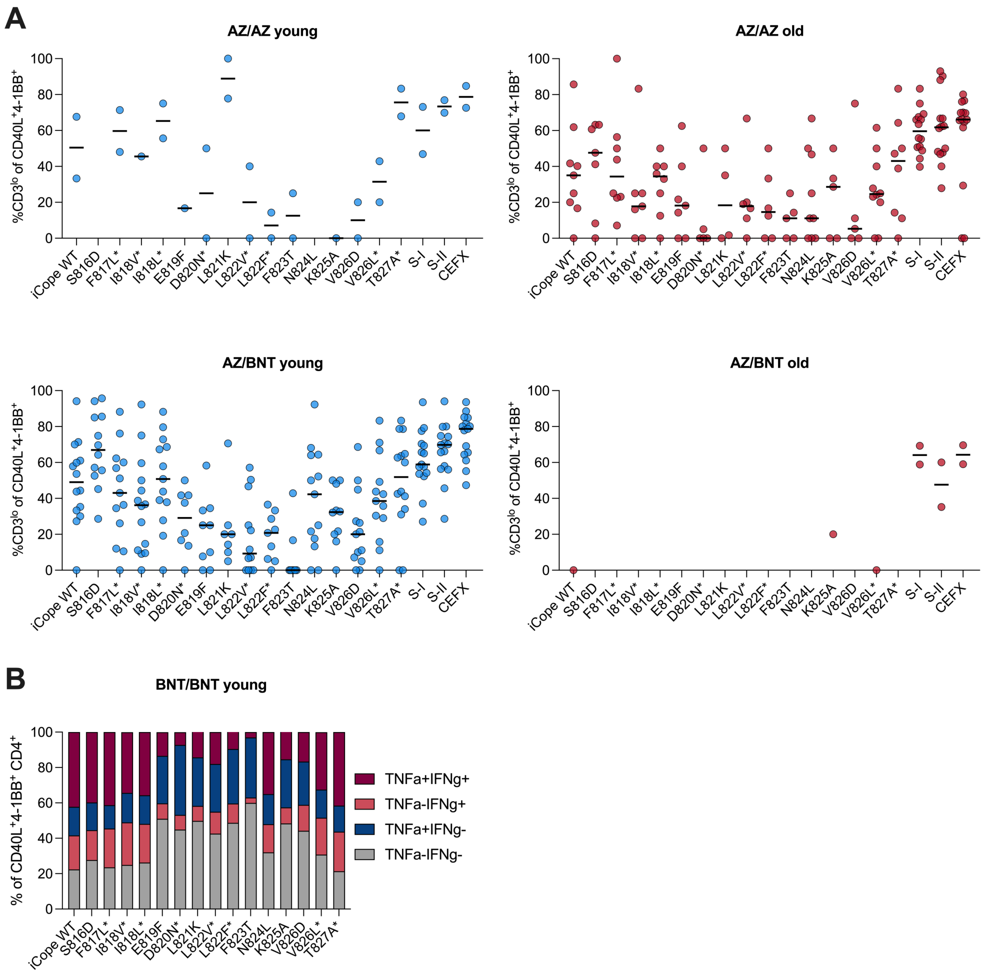


**Fig. S2: $CD3^{lo}$ T cells in AZ/AZ and AZ/BNT vaccinated and mutations affecting the cytokine profile. (A)** Ex vivo stimulation of PBMCs from young and older AZ/AZ (*n*=2/14) and AZ/BNT vaccinated (*n*=16/2) individuals with iCope WT or different mutated iCope peptides and the control pools S-I, S-II and CEFX. Frequencies of $CD3^{lo}$ cells in $CD40L^{+}4\text{-}1BB^{+}$ $CD4^{+}$ T cells are shown for T cell responses with a stim. index ≥1.5. **(B)** Proportion of IFN-γ and/or TNF-α producing T cells among $CD40L^{+}4\text{-}1BB^{+}$ $CD4^{+}$ T cells in the BNT/BNT young after stimulation with iCope WT or the different mutated peptides. * indicates documented iCope mutations.

### Figure S3

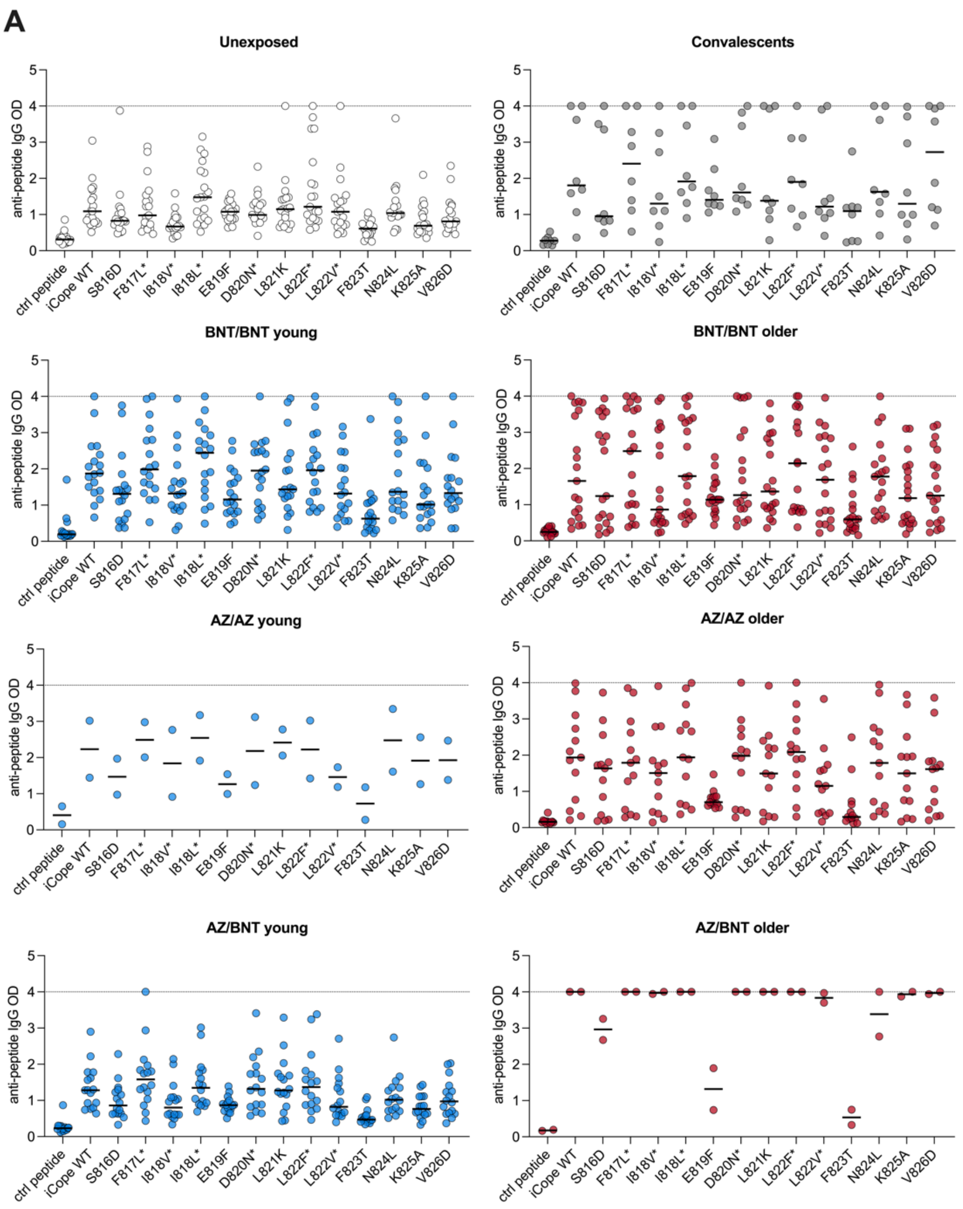


**Fig. S3: Anti-S809-826 humoral immune responses are affected by mutations but not by age. (A)** Optical density of anti-S809-826 peptide IgG (ELISA) relative to indicated mutations. Peptide S1133-1147 was used as internal control (ctrl peptide). Dotted line indicates the upper detection limit. * indicates documented iCope mutations.

### Figure S4

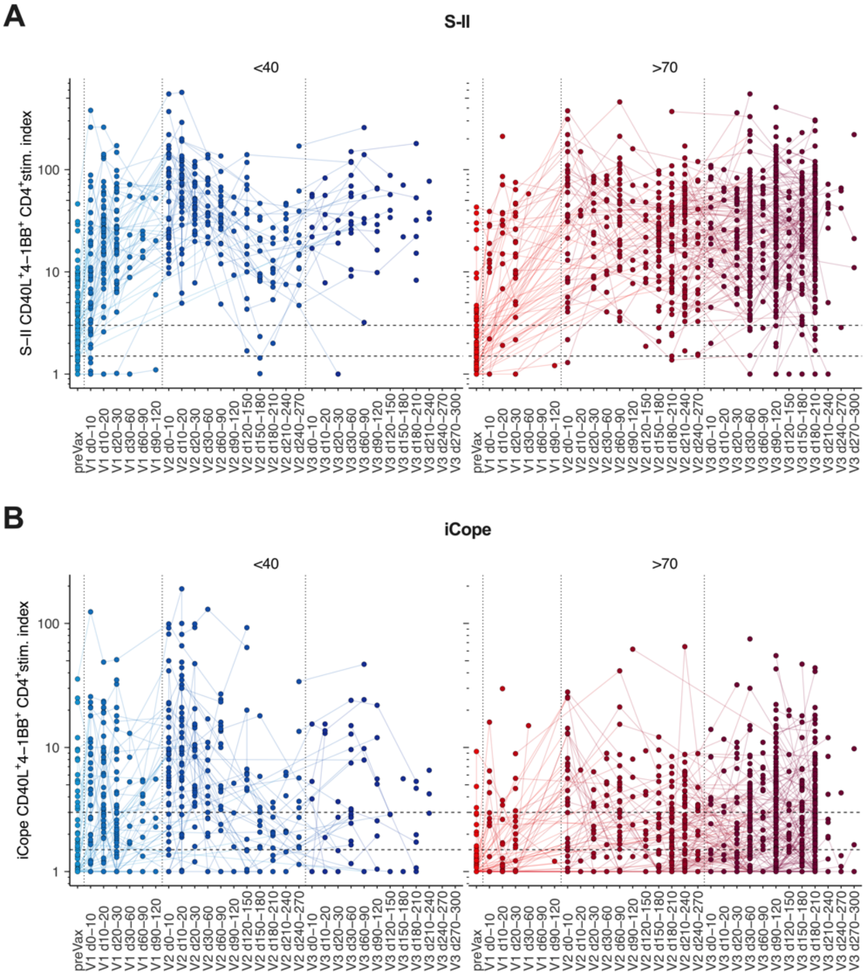


**Fig. S4: S-II and iCope responsiveness over time in young and older. (A)** Ex vivo stimulation of PBMCs from young (<40 years, blue) or older (>70 years, red) individuals with S-II prior to first vaccination (preVax) or at indicated timespans after the first, second or third dose of vaccine (V1-3). **(A, B)** Individual resolution of SI after stimulation with S-II **(A)** or iCope **(B)**. Repeated measurements of the same individuals are connected with lines.

| Cohort | Donor number (#) | Gender (# females) | Mean age and [SD] (years) | Mean time since infection/ 2nd dose [SD] (days) |
|---|---|---|---|---|
| **Unexposed** | 17 | 12 | 26.7 [4.2] | - |
| **Convalescents** | 8 | 7 | 43.6 [12.2] | 286 [57] |
| **BNT/BNT young** | 18 | 11 | 30.0 [7.1] | 24 [6.7] |
| **BNT/BNT older** | 19 | 7 | 76.7 [3.9] | 22 [3.1] |
| **AZ/BNT young** | 16 | 9 | 27.0 [5.8] | 83 [3.6] |
| **AZ/BNT older** | 2 | 1 | 63.0 [4.2] | 75 [12.7] |
| **AZ/AZ young** | 2 | 2 | 30.5 [4.9] | 78 [0.7] |
| **AZ/AZ older** | 14 | 7 | 71.4 [3.4] | 78 [5.9] |

**Table S1: Donor characteristics T cell stimulations and serology.**

| Cohort | Donor number (#) | Gender (#females) | Mean age and [SD] (years) | Time since 3rd dose/infection [SD] (days) |
|---|---|---|---|---|
| **Young vaccinated** | 1-3 | 2 | 29.3 [1.5] | 85.7 [8.1] |
| **Young vaccinated and infected** | 4-9 | 6 | 31.3 [7.3] | 46.1 [33.7] |
| **Older vaccinated** | 10-12 | 2 | 82.3 [1.5] | 80 [12.2] |
| **Older vaccinated and infected** | 13-17 | 1 | 81.0 [3.4] | 54.2 [35.6] |

**Table S2: Donor characteristics single-cell RNA-sequencing.**
